# Thrombo-inflammation analyzed in a validated seven-layer platelet decision model: cellular decisions are tough problems fast and heuristically solved

**DOI:** 10.1101/2024.08.02.606324

**Authors:** Juan Prada, Johannes Balkenhol, Özge Osmanoglu, Maral Afshar, Martin Kaltdorf, Sarah Hofmann, Sebastian von Mammen, Katrin G. Heinze, Harald Schulze, Thomas Dandekar

## Abstract

Decisions in biology happen fast and are driven by evolution to optimize survival chances. In platelets, this is achieved by organizing signaling cascades into rapid decision-funnels with modulatory crosstalk. We show that network decision processes underlying cellular decisions are tough to solve (equivalent to classical satisfiability problems, SAT). Hence, heuristics, modular decision-making, and decision funnels are required for efficient decisions.

We establish this using a seven-layer platelet decision network that agrees well with all available genetic and functional experimental data. Platelet decision cascades are robust to perturbations: For example, receptors such as TRPM7 modulate platelet activity. However, knockouts of the receptors still leave platelets reactive overall. Dynamic control resolves relaying functions from kinases to cytoskeleton alterations. This allows fast execution of platelet shape change or aggregation. Stress conditions can shift platelet decision funnels towards constant activation of aggregation or immune signaling, causing thrombosis or thrombo-inflammation. Based on the network dynamics, we conclude that platelets pragmatically resolve the complex (non-polynomial (NP)) cellular decision problems by using a similar relaxation to those proposed in mathematics – many different configurations end up in similar states. Metamathematical considerations (no mathematical proof) suggest that NP problems are more complex then P problems.

**One sentence abstract:** We show that cellular decision problems like the platelet signaling cascade may need unexpectedly long to solve but in general, they are efficiently solved using heuristics (“decision funnels”), implying fast decisions but the risk of chronic stress and inflammation.

## Introduction

Platelets, the smallest of our blood cells, play a pivotal role in hemostasis and thrombosis, orchestrating a rapid response to vascular injury by forming clots to prevent bleeding [1]. Despite their lack of a nucleus, platelets exhibit complex signaling networks, guiding critical decisions in clot formation and wound healing [2–4]. The rapidity and precision with which platelets activate and aggregate in response to vascular damage [5] show smart and fast decision processes mimicking small computer chips.

In recent years, computational models have increasingly been employed to unravel the intricacies of platelet signaling pathways, offering insights into the dynamic decision-making processes of these cellular fragments [2–5]. These models underscore the need for a robust framework to understand how platelets integrate multiple signals and make decisions in real-time, a process that is remarkably efficient and effective.

Tough decision problems, like decision problems of platelets in complex environments, may need an unexpectedly long time to solve. These problems are called NP problems, which stands for non-deterministically solvable in polynomial time. NP problems (see supplementary introduction) are known for their computational complexity. The significant resources required to reach optimal solutions, finds an intriguing parallel in biological systems. Cells, including platelets, are confronted with myriads of decisions, often needing to solve complex, combinatorial problems under tight time constraints [6].

In this context, platelets emerge as a fascinating model for exploring how biological entities solve complex problems swiftly and effectively. Despite their apparent simplicity, the signaling networks of platelets encompass a level of decisional complexity that could potentially mirror the solving of NP-hard problems in computational systems [7].

This paper aims to analyze the signaling pathways of platelets as a model system, investigating the hypothesis that platelets, through their signaling networks, demonstrate an innate ability to ‘relax’ NP problems into tractable P problems, thus resolving complex decisions with remarkable speed and efficiency. It follows from our results that in the open problem of P vs. NP, nature seems to be on the side of those who claim that NP is NOT equal to P since, after millions of years of evolution, it still has not found a better solution for cellular signaling cascades than a P-relaxation, just as we do.

Using an experimentally well validated *in silico* model of the platelet signaling cascade we first convince the reader that cells have to solve really tough decision problems. We show that they are even equivalent to the famous satisfiability problems in computer science (non-deterministic polynomial problems or NP problems). In computers, this should lead to unexpected stalling of decision processes. However, we can show that in platelet cells this is rarely the case as decision funnels help to find nevertheless heuristic and fast solutions. We support this observation again in the context of published experimental data and evaluating important signaling kinases and modulatory input involved. Moreover, under constant stress there is an inherent risk of changing the system state, for instance, the healthy platelet state can switch to a state of chronic thrombo-inflammatory signaling[8].

As a general conclusion, the tough cellular decision problems are never accurately solved but rather a fast, but not perfect pragmatic cellular response is executed with a risk of changing the healthy system state to unhealthy. Meta-mathematically speaking (no mathematical proof), these biological modelling supports strongly the conjecture that NP problems are always bigger then P (polynomial) problems.

## Material and Methods

### Network reconstruction and dynamic simulations

We extended our previous model of the platelet central cascade [2–4] to include Ca^2+^ and Mg^2+^ transporters, cytoskeletal changes, and environmental feedbacks and we represented the biological context with positive and negative feedbacks. The constructed model is supported by available literature (Table S3). The interactions were classified either as activating (+) or as inhibiting (-). We constructed and visualized the network topology on yED graph editor (https://www.yworks.com/products/yed).

We used Jimena [10] for semiquantitave simulations to represent platelet activation in a biological context. We also reproduced several experimental and pathological conditions by introducing perturbations to our in silico simulations. Details for each simulation can be found in Table S1.

### Analyze platelet signalling: Formal and Boolean Analysis

#### The satisfiability (SAT) problem

SAT is a decision problem that decides whether a Boolean formula, written in the conjunction normal form (CNF), can be evaluated as true. Let *x_i_* be a Boolean variable, which takes only 0 (false) or 1 (true) values. Let *l_i_^j^* be the literal associated with the Boolean variable *x_i_* in the clause *j*. *l_i_^n^* its either *x_i_*, ¬*x_i_* or 0. The literal either takes the value of the Boolean variable or its negation or 0 (false). This last option represents that the Boolean variable is not part of a clause. A clause in the CNF form is a disjunction of literals. A complete Boolean expression is a conjunction of clauses, as seen in **equation S1** (The general form of a Boolean expression in CNF form).

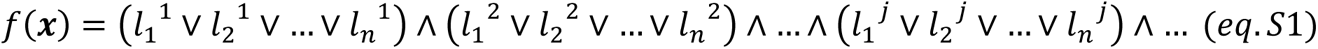

Accordingly, SAT decides whether there exists a set ***x*** of Boolean variables that satisfies (evaluates to true) such an expression f(x). Usually, we set the maximum number of variables allowed in each part of a SAT problem. Because of this, we often call the SAT problem the K-SAT problem. In this name, ‘K’ represents the largest number of variables any part can have. The 2-SAT problem is a P problem, but 3-SAT and all other bigger forms are NP-complete.

Another problem is the Max-SAT problem. It is defined by whether the Boolean expression is satisfiable or not. The idea is to find the combination of Boolean variables ***x*** such that the maximum number of clauses possible is satisfied. These problems are also NP-complete.

### The platelet signaling cascade is translated into a Boolean function

To interpret the platelet signaling cascade as an SAT problem, we first need to transform it into a Boolean network. This transformation is realised simply by assuming that each protein in the network is a Boolean variable and it has only two possible states: 0 (inactive/false) or 1 (active/true).

Now, the edges connecting proteins mean that the protein on the ending side depends on the protein on the starting side. This means that it can be activated or inhibited by the parent protein of the edge depending on the shape of the edge termination. An arrow-shaped edge is used for activation, and an orthogonal short line is used for inhibition. For example, STIM1 activates ORAI1, which in Boolean terms would be written as *STIM*1 = *ORAI*1. As an inhibitory example, see that P2Y12 inhibits AC. In this case, the Boolean expression would be *AC* = ¬*P*2*Y*12.

If multiple edges arrive at a protein, it means it is controlled by these respective incoming signals. Overall, this yields a logical OR among all the parent proteins. For example, the Boolean formula for the IP3R node in **Fig. 2** is:

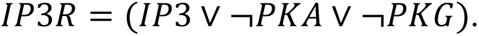

Here, the usage of OR is sufficient for the modeling [11]: In cellular signaling, it is common to encounter redundant activation pathways. Essentially, this operates much like an ‘OR’ gate, where multiple routes can independently lead to the activation of the same function or response. Moreover, the reaction of multiple molecules at once is rare. Usually, compounds are formed in a sequential form. If one wants to represent an AND, the usual trick is to create an artificial intermediate inhibitory node. As a result, the PPI network can be transcribed into a Boolean expression using this simple convention.

### Platelet cellular decisions formally represent a satisfiability (SAT) problem

As outlined above, several processes must be initiated to have a successful clot formation process that prevents the organism from bleeding. We included 5 of those processes in the network: Granule release (GR), Aggregation (Ag), Stress fiber formation (SFF), Lamellipodia (La), and Shape change (SC). We refer to the coagulation process as CP. The full CP requires each of the five processes. The Boolean expression corresponding to that situation would be as follows:

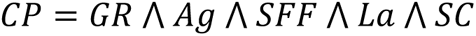

We can further expand this Boolean expression by replacing each of the processes involved with the corresponding set of activating parent nodes. In that case, the coagulation process would be described as follows:

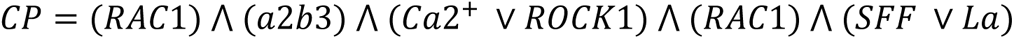

Following the procedure of replacing the nodes with their Boolean activation equation, we would obtain:

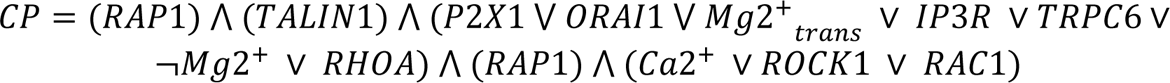

Then, recursively following the same procedure, we obtain:

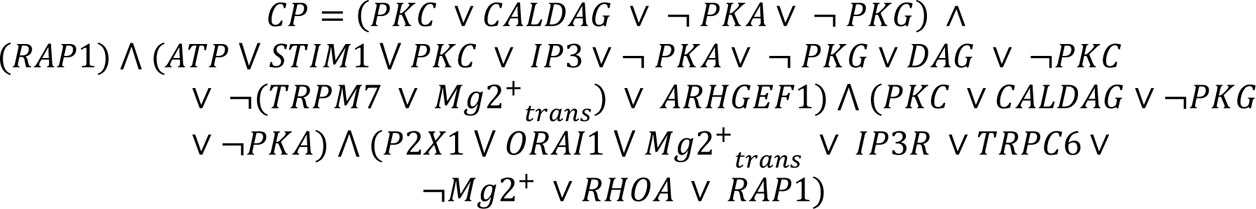

Finally, by solving the negated OR, we get:

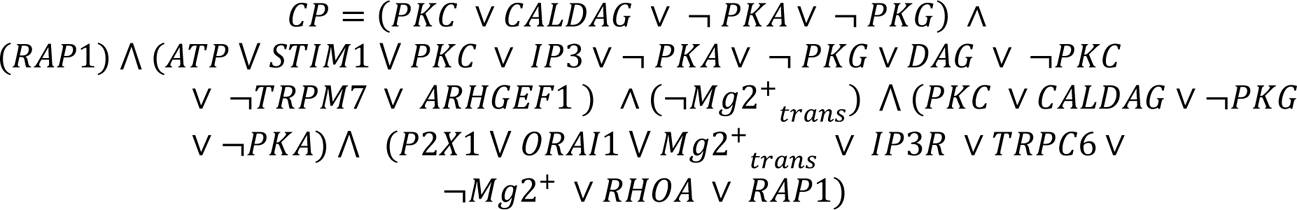

Notice that this expression is not final and that we could go on replacing different nodes with their more complex, respective activating function, but there is no need to go further. The expression we observe is already a clear example of a Boolean expression in the CNF (conjunctive normal) form.

Observing this expression allows us to formulate the platelet signaling cascade problem as an NP problem, precisely the MaxSAT problem (MaxSAT: satisfy as many conditions as possible). The idea is that to perform the coagulation process, the organism needs to activate each of the processes involved (Aggregation, Stress Fiber Formation, Lamellipodia, Shape Change). It does not necessarily need to activate all of them simultaneously, but it should activate as many of them as possible since bleeding can be deathly very quickly. Hence, we assume that the organism is trying to find the combination of active and nonactive proteins which achieves the maximum number of active coagulation processes. In the best-case scenario, all of them are active at once. This is the exact definition of the MaxSAT problem, as explained above. Accordingly, we are trying to find the Boolean variables (proteins) combination that maximizes the number of satisfied clauses (coagulation processes).

In the same way, we proceeded with the platelet signaling cascade, it is possible to interpret any PPI network as a MaxSAT NP problem. Nevertheless, as stated before, our organism deals with such issues at any moment and seems to solve them accurately and quickly. We will now try to understand how this happens and what we can learn from it.

### Integer program formulation of MaxSat (IP)

The MaxSat solver follows this formalism:

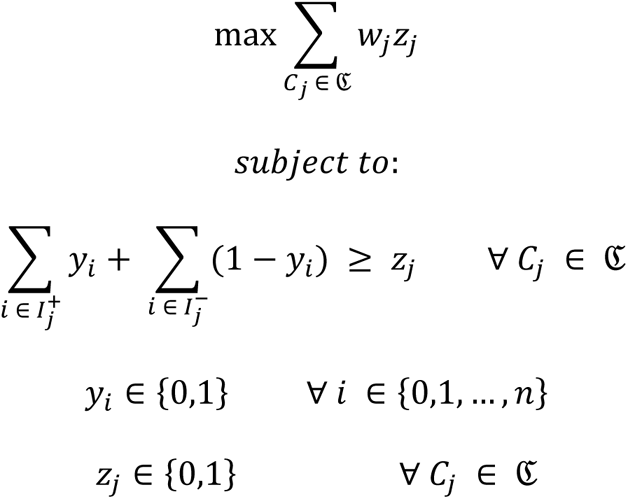

*C_j_* is the clause *j* in the set of all clauses ℭ. Where, *w_j_* is the weight assigned to clause *j*. *z_j_* is 1 if the clause *j* is satisfied and 0 otherwise. *y_i_* is a literal in the clause which equals 1 if the variable *x_i_* is true and 0 if *x_i_* is false. The objective function is to maximize the value obtained from satisfying clauses with their respective assigned weight. If we set all weights to 1, the program tries to satisfy as many clauses as possible. The first constraint ensures that at least one variable is positive or a negated variable is negative because the clauses are conjunctions of variables, so one *true* value is enough to satisfy the clause. The second restriction assures the Boolean nature of the variables. The third condition ensures that the clauses will be evaluated as *True* or *False*.

### Computational Analysis of Biological Networks using SAT and MAX-SAT Solvers

In this study, we employed SAT and MAX-SAT solvers to model and analyze the logical structure of the platelet network. Specifically:

- **SAT Solvers:** These were used to determine the exact conditions under which the platelet network is activated. This involves identifying scenarios where the Boolean formula representing the network is fully satisfied.
- **MAX-SAT Solvers:** These solvers provided insights into the network’s behavior when not all conditions can be simultaneously met. They helped to highlight the most critical components and pathways by identifying the maximum number of clauses that can be satisfied concurrently.

**Heuristic and Decision Funnel Analysis -** We conducted a comparative analysis of the solutions obtained from SAT and MAX-SAT solvers to:

- Identify the heuristics and decision funnels employed by the platelet network.
- Understand the network’s preferred pathways, critical nodes, and potential error-prone routes, offering insights into the network’s decision-making processes under various physiological and pathophysiological conditions.

**Applications in Systems Biology -** Our methodology was applied in several key areas within systems biology:

- **Critical Node Identification:** We pinpointed essential components within the platelet network, assessing the impact of the presence or absence of these components on the overall functionality of the network.
- **Pathway Redundancy and Robustness:** Our approach helped reveal redundant pathways and their contributions to the network’s robustness, enhancing our understanding of how biological systems maintain functionality under stress or perturbations.
- **Error Analysis and Pathophysiological Insights:** We identified error-prone pathways and preferred states within the network, providing crucial insights into potential pathophysiological mechanisms. This analysis is instrumental in guiding future experimental and therapeutic strategies.

More details on all methods are found in the supplementary material.

## Results

### An experimentally validated platelet signalling network to analyse cellular decision problems

This study analyzes how platelets solve complex problems quickly by translating the platelet protein-protein interaction (PPI) network into a Boolean satisfiability (SAT) problem. The SAT problem is NP-complete and outlined in the Material and Methods section. K-SAT and Max-SAT problems are found in the PPI networks in living organisms. Any directed PPI network can be interpreted as a Boolean network. Therefore, Boolean expressions can be extracted from it. Here, the SAT problem is formulated for the platelet central signaling network previously defined by systems biological modeling and is extended here [2–4, 12]. **Figure 1** shows our model for platelet signaling. This signaling cascade is responsible for forming a platelet plug in the bloodstream to stop bleeding when an injury occurs.

**Figure 1:**
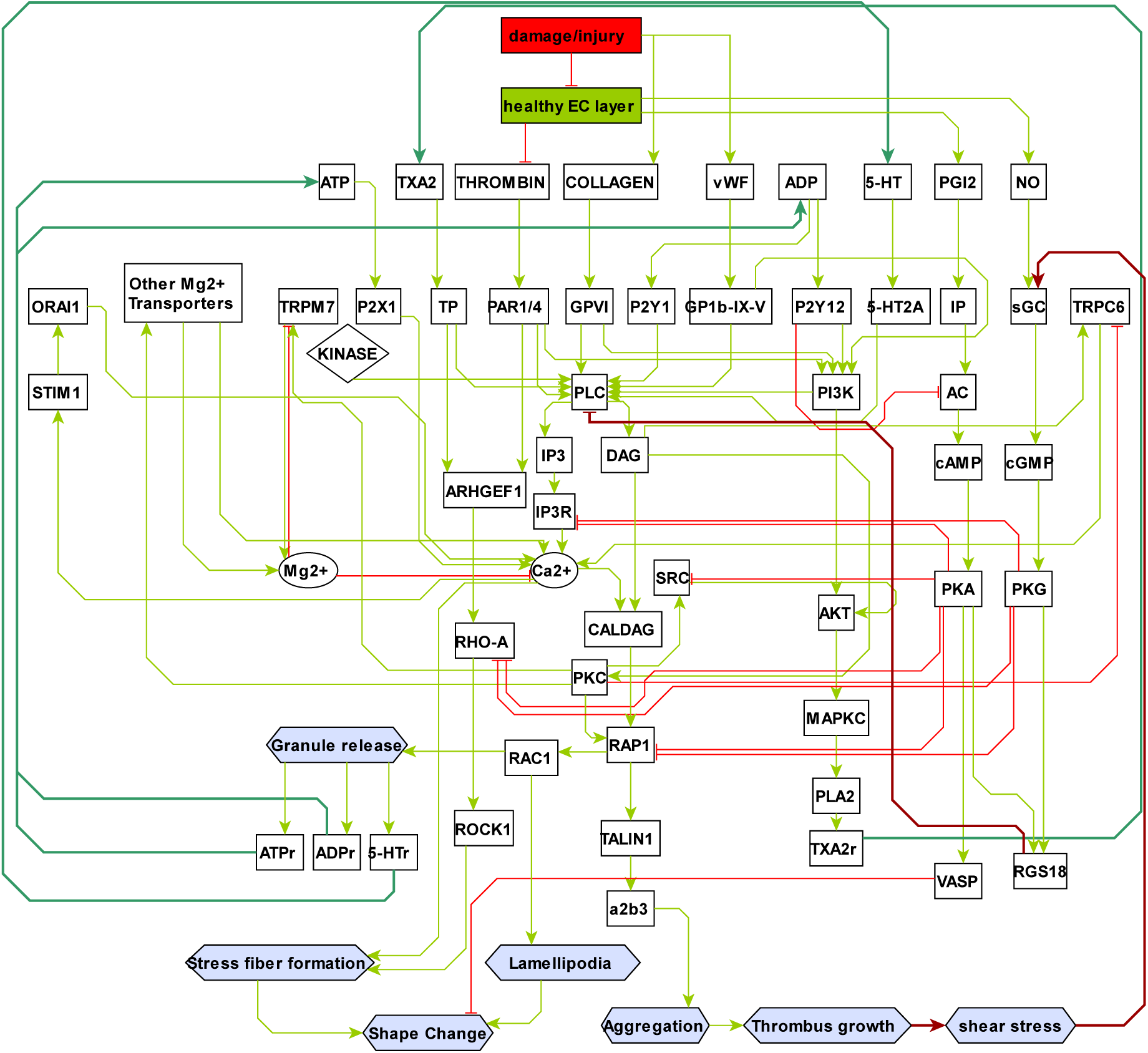
Cell signaling model of Platelet activation. White boxes represent proteins. Their input from other proteins by interactions is shown as activating (green arrows) or inhibitory (red blunted arrows). The resulting cellular functions are shown as violet hexagons (e.g. shape change). Activating interactions in signal amplification are shown in thick green arrows (positive feedback), while internal and external brakes are shown in thick red arrows (negative feedback). An electronic version of the network is added in the supplementary materials (SupplementaryFile2.graphml).

The central signaling cascade represents major players in platelet signaling, such as proteins, chemicals, and behaviors as nodes and their relationship as directed interactions. The interconnections, including activation inhibition, have been optimized according to recently available experimental studies and data (Table S1). This is an updated model of the central signaling cascade of the platelet, which incorporates all modulatory crosstalk and agrees with experimental data available until today. There are several features that are critical for platelet function, as observed in the model and in experiments:

1. Calcium plays a pivotal role in the platelet activation cascade, participating in processes such as granule release, stress fiber formation, lamellipodia extension, and shape change.
2. Protein kinases PKA and PKG inhibit platelet activation by targeting RAP1, RHOA, IP3R, and SRC and activating VASP and RGS18.
3. Shape change requires formation of stress fibers and lamellipodia but inactivation of VASP.
4. Active PKC facilitates signal transmission to SRC, TRPM7, and Mg^2+^ transporters. However, inactivation of PKC is essential for TRPC6-mediated Ca^2+^ transport. Throughout its connectivity, the signaling model represents in detail the antagonistic roles between Ca^2+^ and Mg^2+^ in platelets.
5. PLC serves as a converging point for numerous signaling pathways activated downstream of platelet surgace receptors.

### Dynamic simulations: Analyze platelet problem-solving by analyzing dynamic simulations

#### Platelet inhibition

Figure 2 presents the simulation of the platelet protein-protein interaction network’s quiescent (non-active) state, serving as the reference state for subsequent dynamic simulations. In Figure 2A, the quiescent state is visualized in the network, delineating both active and inactive pathways. The network architecture integrates two input node layers: environmental stimuli, such as damage to endothelial cell (EC) layers, and platelet receptors that detect environmental changes. Internal processing layers assimilate this external information, while output layers act as effectors of behavior, manifesting in platelet shape change and aggregation. A feedback loop from the behavior nodes to the external input nodes indicates environmental alterations triggered by changes in platelet behavior.

**Figure 2:**
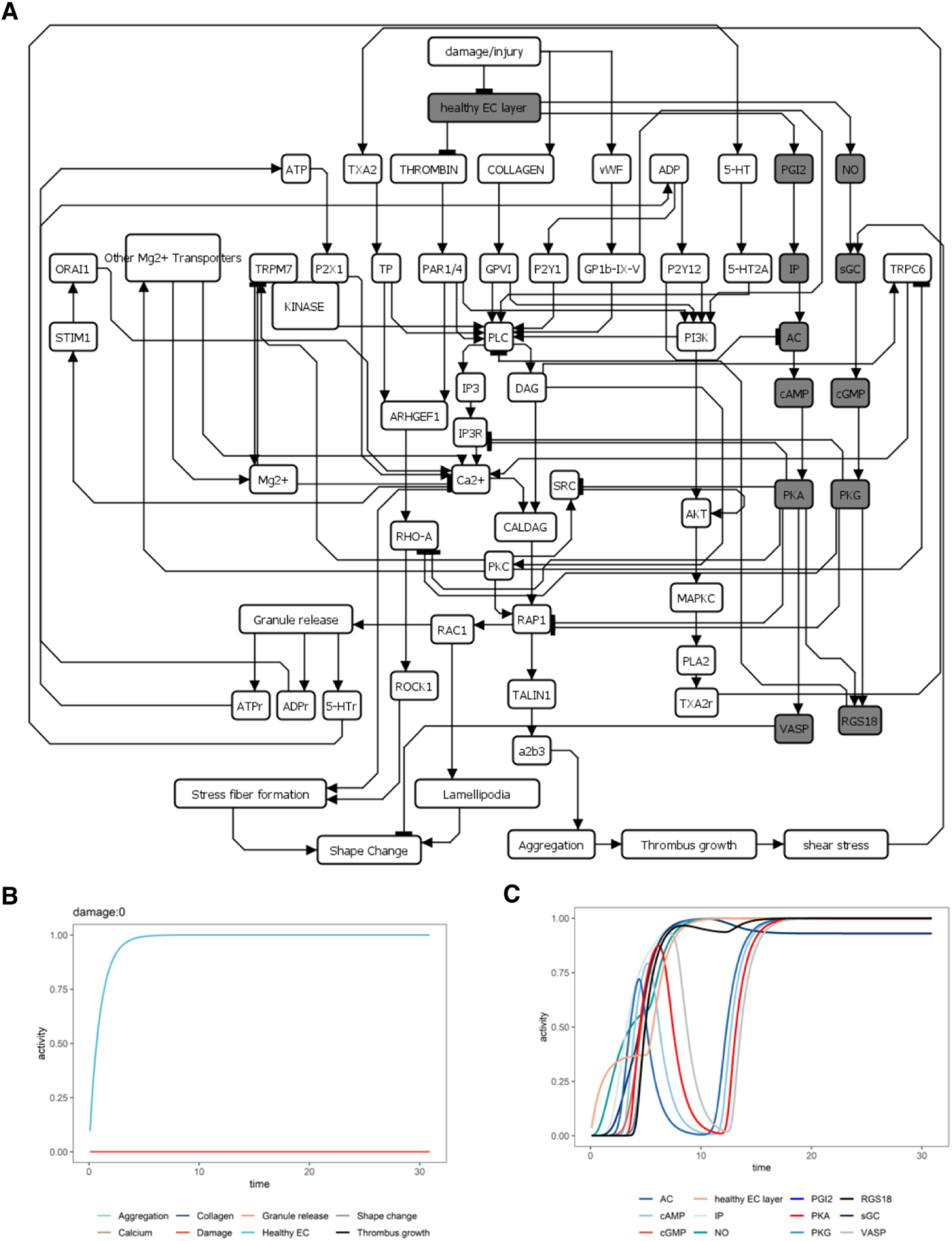
Quiescent state simulation. (A) Indicated is the platelet network of Fig. 2 and the activation of the nodes between values of 0 and 1 (white: no activation; grey: high activation). The network includes two input node layers: environmental stimuli such as damage to EC layers and platelet receptors (sensors that sense environmental changes). Further, internal processing layers process the external information and output layers that are effectors of behavior, e.g., shape change and aggregation. A feedback loop from behavior nodes to external input nodes represents changes in the environment triggered by platelet behavior changes. The activation of the nodes changes over time. Thus, critical nodes can be monitored, indicating node and network behavior in a reference quiescent state maintained by external PGI2 release(B) or by thrombomodulin release (C).

Without an injury, the endothelium is a non-adhesive layer that does not promote platelet activation. In other words, platelets are in a quiescent state. When the platelet simulations are executed without any external input (i.e., in the absence of damage), the platelet signaling is inactive (Fig. 2A). Platelet inhibitory signaling pathways like cGMP, cAMP signaling, and RGS18 are fully active as self-inhibitory mechanisms and keep platelet activation from happening. (Fig. 2A). This is achieved by endothelial cells via the secretion of PGI2 (Prostaglandin I2) and NO (nitric oxide) (Fig. 2A-B). Both activate platelet inhibitory signaling via cyclic nucleotides (cAMP, cGMP; Fig. 2C and also shown by [13]). Another way that ECs keep platelets quiescent is via thrombomodulin, which inactivates thrombin (Fig. 2C and also shown by [13]). Our model represents this by thrombin that inhibits the healthy EC layer (Endothelial Cell layer).

#### Platelet activation

In contrast to the quiescent state, platelet activation is rapidly initiated upon injury (Fig. 3). This transition is marked by the exposure of subendothelial collagen and the interruption of platelet inhibitory signaling from the healthy EC layer, persisting until the damage is repaired. The initiation of platelet activation processes, such as calcium signaling, granule release, shape change, and platelet aggregation, is represented in Figure 3 (blue). These processes augment the size of the thrombus at the injury site, with shear pressure on the thrombus escalating in proportion to its size (Fig. 3; black). However, this activation is not perpetual. Upon attaining the necessary size for damage repair, the healthy EC layer is reconstituted (red), culminating in reinstating internal like RGS18’s inhibition of G protein signaling and external platelet inhibition mechanisms such as mechanosensitive cGMP signaling.

**Figure 3:**
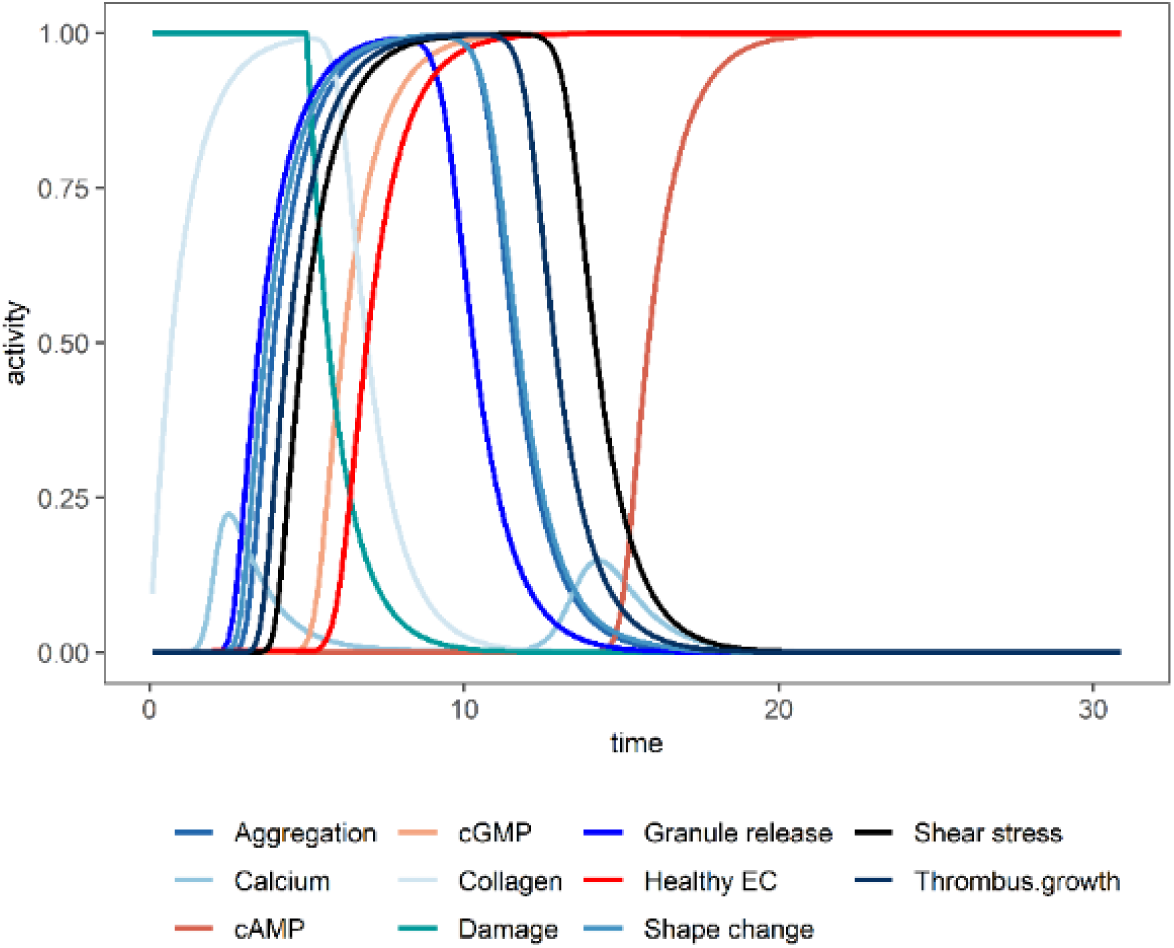
Damage to the EC layer immediately leads to collagen activation and platelet activation. Indicated here are the activation values over time of platelet activation markers (in blues), platelet aggregation processes (black), and EC layer formation (red). Here, platelets are activated by setting the value of the node damage to 1 for 5 time units.

#### Platelet decision funnels: Robustness to receptor modifications due to signal amplification in a feedback loop (decision funnel in time)

Initially, platelet activation is triggered by collagen binding to receptors at the injury site, with subsequent adhesion and downstream signaling leading to shape changes, granule release, and thromboxane generation (Fig. 3). This activation is further amplified through GPCR activation and signal transduction events, culminating in platelet aggregation facilitated by αIIbβ3 interactions with fibrinogen or vWF. The system’s robustness is underscored by a positive feedback loop (external control) ensuring rapid and robust platelet aggregation, as evidenced in our model’s resistance to TRPM7 knockout (KO) (Fig. 4). It is shown that platelet reactivity is dependent on damage (1 to 0.1), but the TRPM7 KO has no significant effect on the platelet activation decision.

**Figure 4:**
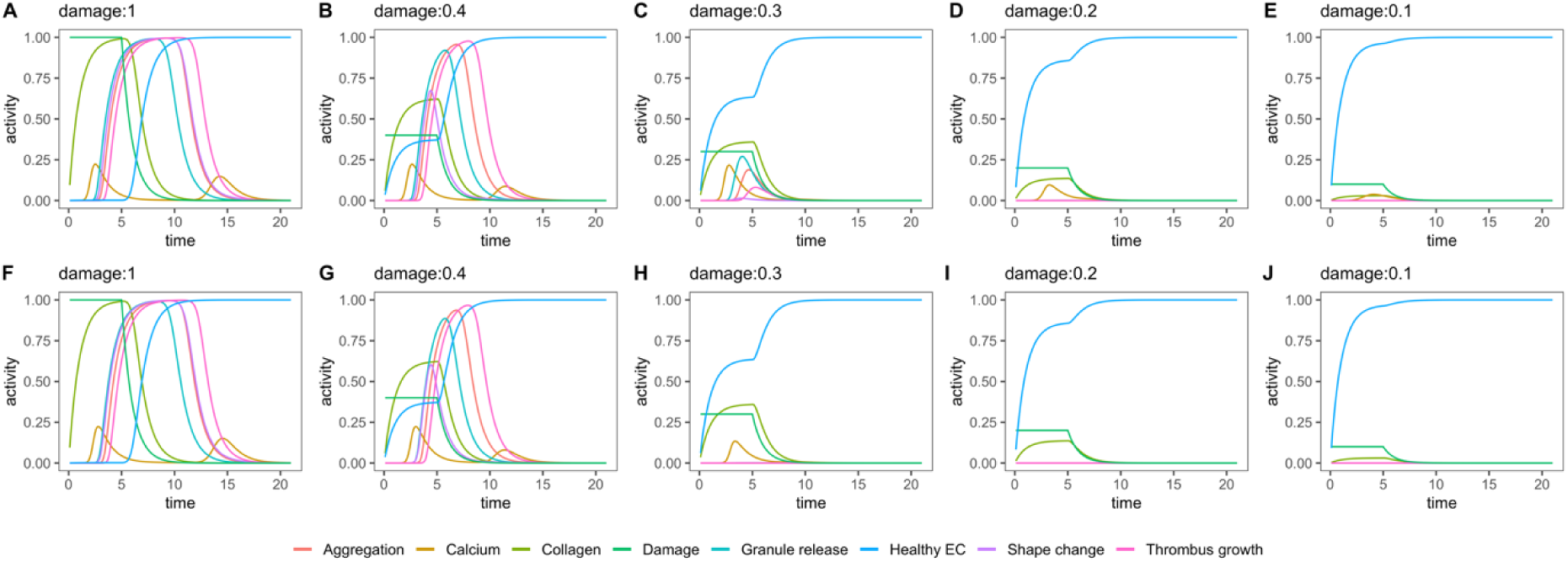
TRMP7 knockout (TRPM7-/-) simulation with external control layers. TRPM7 Kinase domain is set as constitutively active in wild-type (WT, up: A-E), and 0 in TRPM7 knockout (KO, down: F-J). Different damage input values stimulate platelet signaling in both conditions. Threshold analysis determined the intermediate value of collagen to be 0.45 for WT and 0.45 for TRPM7^-/-^ (defined as the minimum collagen stimulus required to reach a full irreversible aggregation (value 1)) and the lower value to be 0.35 for WT and 0.4 for TRPM7^-/-^ (the minimum collagen stimulus to activate the irreversible aggregation to a higher value than the collagen input).

Moreover, the critical role of external control layers in stabilizing decision-making processes or funnel points is starkly evident when examining damage response in a network devoid of such external modulation. The influence of knockout (KO) events is markedly accentuated in scenarios lacking signal amplification or the nuanced context provided by the biological microenvironment, as depicted in Figures 5 and **S1-2**. In these scenarios, certain levels of damage still permit platelet reactivity (e.g., between 0.4 to 0.2; see **Figures 5 A, C, E, G**).

**Figure 5:**
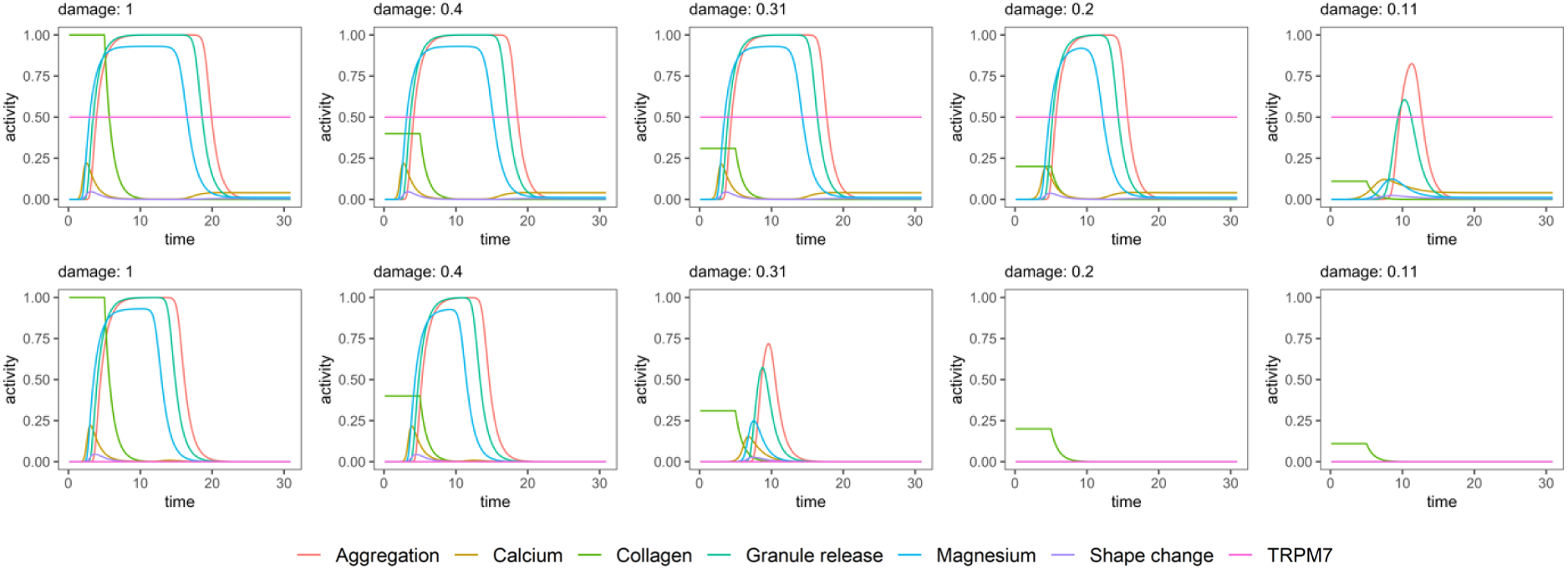
TRMP7 knockout (TRPM7-/-) threshold analysis and simulation without external control layers. Stimulation of the platelet signaling network with different levels of collagen input. TRPM7 Kinase domain is set as constitutively active in wild-type (WT, upper row), and 0 in TRPM7 knockout (-/-, lower row). Threshold analysis determined the intermediate value of collagen to be 0.2 for WT and 0.4 for TRPM7-/- (defined as the minimum collagen stimulus required to reach a full irreversible aggregation (value 1)) and the lower value to be 0.11 for WT and 0.31 for TRPM7-/- (the minimum collagen stimulus to activate the irreversible aggregation to a higher value than the collagen input).

Nevertheless, these networks exhibit heightened sensitivity to TRPM7 KO, highlighting the decreased damage response under conditions where external stabilizing influences are absent (as illustrated in **Figures 5B, D, F, H**).

#### A shift of decision funnels in the platelet is causing chronic inflammation

In thrombo-inflammation, the loss of EC control over thrombosis and inflammation results in heightened platelet activity. ECs normally exert anti-thrombotic and anti-inflammatory influences. However, during thrombo-inflammation, the balance between pro- and anticoagulant properties of the endothelium can be disrupted, leading to sustained platelet activation and immune cell recruitment. In the absence of EC control, platelets become hyperactive and continuously interact with the endothelium and immune cells [14, 15] . Our simulations demonstrated this, showing a cyclic pattern of platelet activation in the persistent presence of procoagulant stimuli or when EC control is compromised (**Fig. S3-4).**

Next, we analyze different mechanisms of how this shift in balance happens and how the resulting chronic inflammation sustains platelet activation and increases the risk of thrombotic events and inflammation. We show this shift after integrating the mechanisms behind inflammation caused by atherosclerosis, inflammatory arthritis (IA), and in aging in our model (**Fig. S5**) which confirms the effects of chronic inflammation stimulating platelet aggregation (**Fig. 6 and S5-6**).

**Figure 6:**
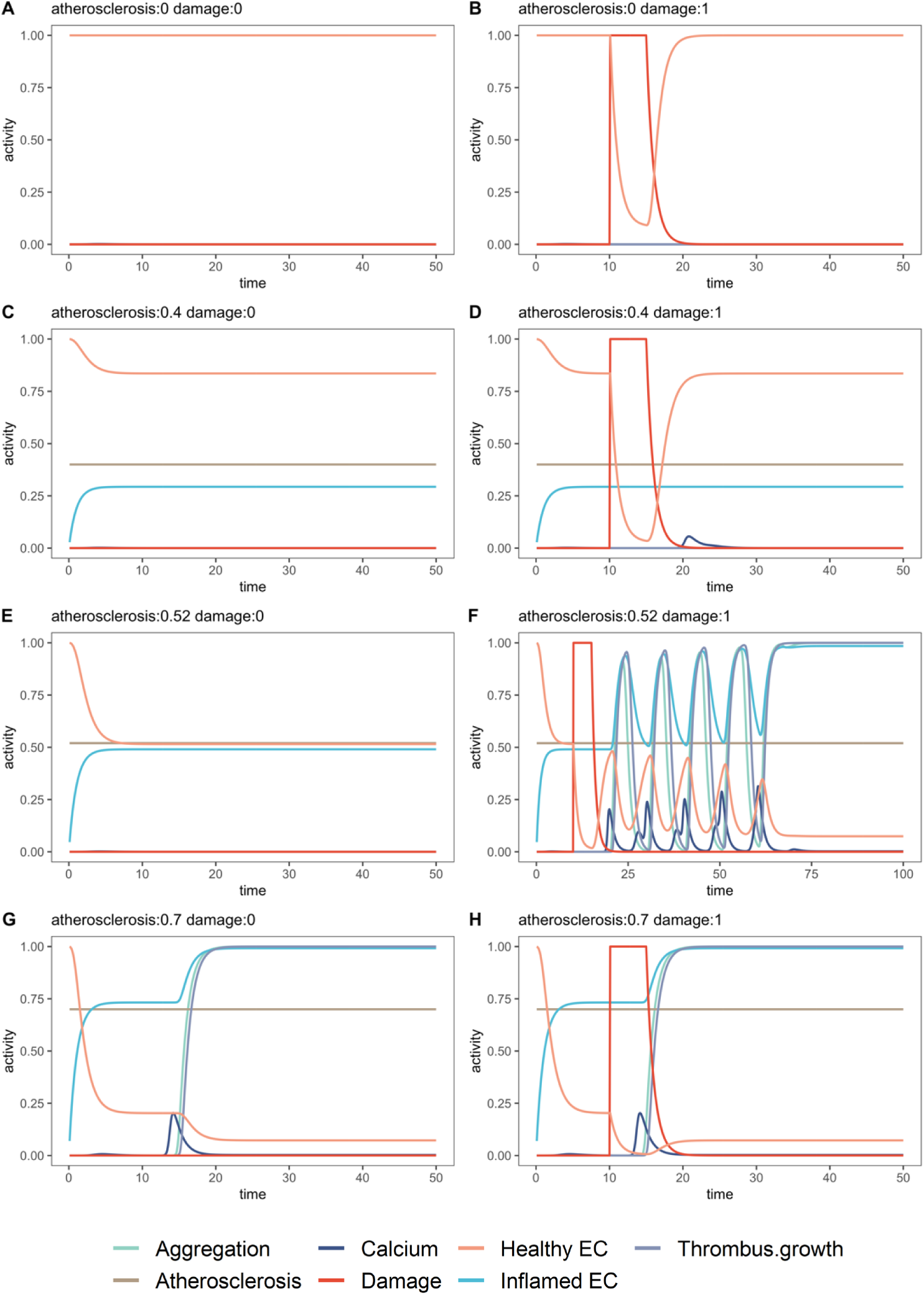
Atherosclerosis simulation with damage input (right) and without damage (left). Values below 0.52 for atherosclerosis do not lead to any aberrations in platelet aggregation. Values between 0.52 and 0.7 only lead to aberrant aggregation when there is a damaged input. Values equal to and bigger than 0.7 lead to aberrant aggregation even though no damage is present.

The simulation of atherosclerosis scenarios revealed that atherothrombosis occurred at high inflammation levels (atherosclerosis ≥ 0.7) in the absence of damage. Medium inflammation levels (0.52≤atherosclerosis<0.7) led to aberrant platelet aggregation only in the presence of damage. However, low-grade inflammation (atherosclerosis < 0.52) did not result in atherothrombosis, even with a damage stimulus, preserving a healthy EC layer and its anti-thrombotic profile (Fig. 6).

The model integrating IA-induced pro-thrombotic EC profiles and platelet hyperactivation (**Fig. S5**) showed a stronger effect on platelet aggregation compared to atherosclerosis. Even at lower/medium IA levels (IA≥0.58), full activation of platelet aggregation occurred, even in the absence of damage input (**Fig. S6**).

In our model, we can also show how aging shifts the signaling of platelet receptors Gp1bα and GPIX, as well as inflammation through TNF-α levels. Although metabolic reprogramming was not considered in our model, our simulations closely mirrored the results seen in platelet hyperactivation in IA-induced inflammation (**Fig. S7**).

### Analyze platelet problem-solving by formulating it as satisfyability (SAT) problems

#### Formulate the platelet network as a Boolean SAT problem

The signaling cascade of proteins is represented as an SAT (Boolean satisfiability) problem by transforming it into a Boolean network to analyze the SAT equivalent of platelet decisions. Proteins are treated as Boolean variables with two states (active or inactive), and their interactions (activation or inhibition) are depicted through specific edge shapes in the network. Complex relationships where multiple proteins influence a single protein are captured using logical operators like OR, representing redundant pathways. This approach is then applied to model the coagulation process (CP), breaking it down into five critical subprocesses, each represented by a Boolean expression involving various activating parent nodes. These expressions are further refined by recursively replacing nodes with their respective Boolean equations, ultimately leading to a comprehensive representation of the CP as a complex interplay of numerous proteins and pathways (**equation 1**). Detailed methodology and the stepwise transformation of the signaling cascade into a Boolean network are elaborated in the **Material and Methods** section of the study.

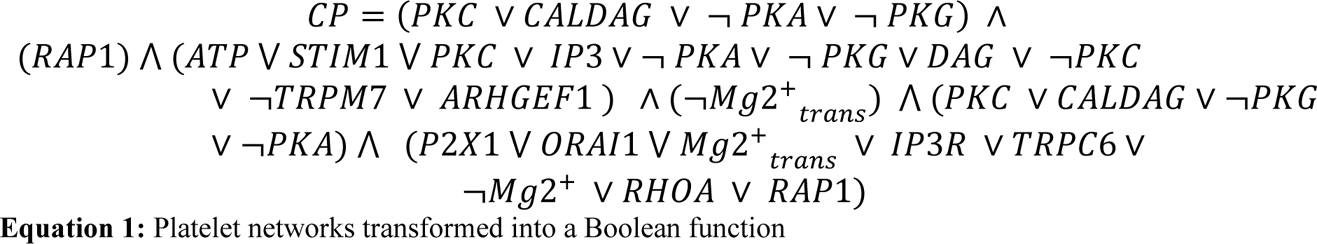

#### Cellular decisions present a MaxSat problem which can be solved by optimization

The MaxSat problem is often presented as a maximization problem, which facilitates its analysis and allows for some estimations of the expected error. We will also present the problem in a way that allows us to relate it to some biological aspects. This formulation is used in many articles. We closely follow the one presented in Goemans Williamson [16].

##### MaxSat relaxation: Theoretical consideration on the organic solution of MaxSAT

In mathematics, when dealing with categorical decision problems like MaxSAT, a standard first step to relax the problem from NP to P is to loosen the categorical nature of the variables and allow them to have a continuous spectrum of values. In the case of Boolean variables, this would mean that we allow the variables to have a real value between 0 and 1 instead of the binary value of 0 or 1 (**see M&M Integer program formulation of MaxSat (IP)**). This relaxation occurs naturally in the case of PPI networks since each protein node in the network represents not a single entity but a protein concentration. Concentration values are not either present or absent but correspond to a continuous variable. The optimization problem stated before (IP) in its relaxed version becomes a linear program.

##### MaxSat relaxation: Linear program formulation of MaxSat (LP)

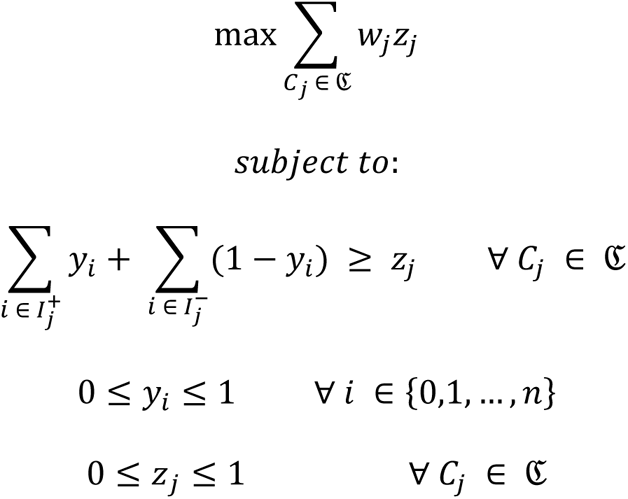

Notice that the only difference is that now the variables *y* and *z* become continuous in the interval from 0 to 1. We see here a clear conclusion: the PPI network problem becomes NP only in our mathematically accurate representation, but in the organism, for instance, in platelets, it remains in P. Evolutionary selection has allowed for efficient adaptation in real life. The fact that the problem is naturally in P explains that the body manages to solve it so fast. Being in P and restricting the decision to a specific set of proteins (some hundreds in most cases) means the organism can find a feasible solution quite fast. Now, the question remains: How can the solution be so accurate? In computation, a strategy that seeks to improve a solution is called a heuristic. We want to interpret the heuristics proposed by Goemans and Williams [16] in the light of living organisms and see whether there could be a relation between them or find which heuristics living organisms rely on in the protein signaling cascade resolution. A commonly used heuristic is to solve the linear program and to use the value of *y_i_* as a probability to decide whether *x_i_* is *true* or *false.* This simple heuristic achieves a 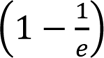 -approximation.

This means that there is a guarantee that the solution to the relaxed problem is at least within the 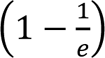 range from the optimal solution of the NP problem. In the Goemans Williams paper [16]. they proposed a 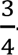-approximation, which resembles the previous heuristic but with a slight modification. The idea is not to use the value of *y_i_* directly as the probability of *x_i_* but to create a function such that:

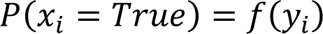

To this end, not just any function can be used, but the ones in a set of functions which have the so-called 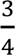 property, i.e., all functions in the following set as defined by Goemans Williamson [16]:

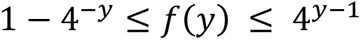

Hence, any function contained between these two functions in the interval from 0 to 1 could be used to randomly assign the values of the *x_i_* variables. A visual representation of such a space is presented in Figure 7.

**Figure 7:**
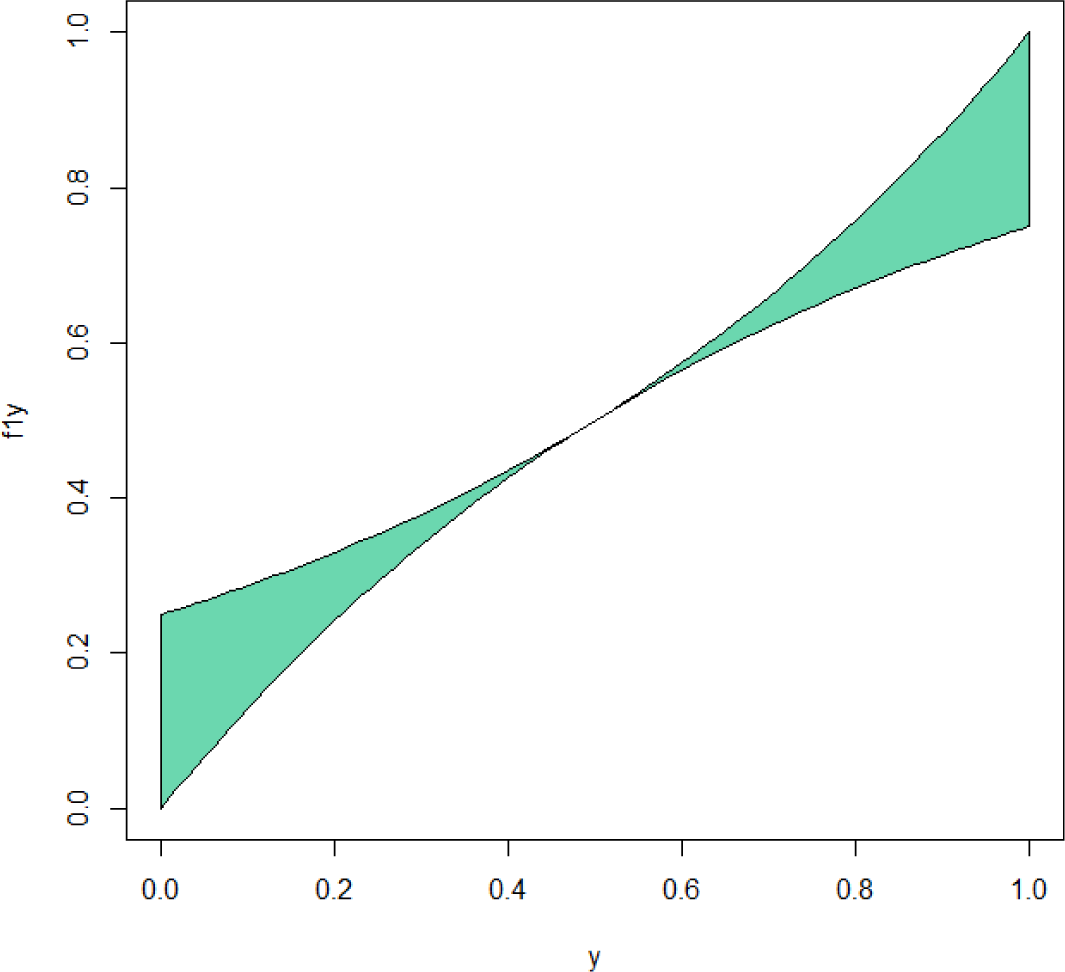
The set of ¾ functions apply to biological solutions for complex problems. The green area represents the space where the 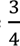 functions have their solution.

#### Computational Analysis of the Platelet decision network using SAT and MAX-SAT Solver

In our analysis, we employed SAT and MAX-SAT solvers to delve into the complex dynamics of a Boolean network representing platelet signaling pathways. This approach has uncovered pivotal insights into the network’s operational principles and hierarchies.

The network analysis revealed various pathway categories: Critical Nodes as essential for functionality, highlighting their pivotal role in system activation and integrity-dispensable nodes or pathways, which don’t significantly affect outcomes, contribute to the network’s flexibility and fault tolerance. Redundant Pathways enhance the system’s robustness by preventing reliance on single components. Preferred Pathways indicate the system’s efficiency-driven choices, while Error-Prone Pathways highlight potential inaccuracies or inefficiencies, offering insights into vulnerabilities. Decision Funnels, as crucial convergence points for signals, are essential in understanding the network’s regulatory mechanisms. Lastly, the comparison between “Ideal vs. Realistic Solutions” provides a nuanced view of the system’s operational heuristics, revealing its priorities, compromises, and patterns, offering valuable insights for system understanding and potential improvement.

Our simulation of the platelet activation network via SAT solvers demonstrated that various combinations of signaling inputs can lead to platelet activation, signaling the onset of aggregation and clotting. This plurality of pathways underscores the biological system’s complexity and redundancy. However, the indispensability of elements like RAP1 and Mg^2+^transport for activation signals our attention to their critical roles in the biological context. These insights, summarized in **Table 1**, underscore the intricate interplay of heuristics, decision-making, and redundancy in the cellular signaling network, offering a rich tapestry of information for understanding and optimizing the network’s behavior in biological systems.

**Table 1:** Computational analysis options of biological networks using SAT and MaxSat solver^1^.

| Category | Description | Example Components/Pathways |
| --- | --- | --- |
| Critical Nodes | Components that are consistently present in MAX-SAT solutions, indicating they are essential for activation. | PKC, CALDAG, ... |
| Dispensable Nodes/Pathways | Components that can be absent without significantly affecting the overall activation of the network. | TRPM7, PKA (when other activating pathways are active), ... |
| Redundant Pathways | Components or pathways that provide overlapping functionality, offering robustness to the system. | PKC and CALDAG (if both lead to similar activation patterns), ... |
| Preferred Pathways | Pathways that are more frequently used or favored in the network's activation process. | PKC → IP3 → RAP1, ... |
| Error-Prone Pathways | Pathways that often lead to suboptimal activation or are associated with system malfunctions. | Pathways involving $\neg Mg^{2+}$ when other critical nodes are inactive, ... |
| Decision Funnels | Points in the network where multiple pathways converge, leading to a key decision or activation. And number/percentage of possible configuration of iterals leadint o the same result | RAP1 as a convergence point for multiple upstream signals, ... |

| Category | Description | Example<br>Components/Pathways |
| --- | --- | --- |
| Ideal vs. Realistic<br>Solutions | Comparing SAT (ideal) and MAX-SAT (realistic) solutions to highlight the compromises made by the system. | The presence/absence of certain nodes in SAT vs. MAX-SAT solutions. |

<sup>1</sup>This allows to examine in a mathematical way the decision heuristics used by platelets captured in our seven layered decision model agreeing well with the experimental data.

## Discussion

This computational heuristic analysis represents a significant advancement in modelling platelet signaling networks, a seven-layered protein network model of involved decision processes agreeing well with all experimental data on platelets considered in its construction. Beyond the example we offer an analytical approach for a comprehensive and nuanced understanding of further complex biological networks. By bridging computational logic with systems biology, our methodology not only elucidates the operational principles of platelet networks but also sets a precedent for analyzing other biological systems with similar complexity. We show here for the first time that cellular decision networks are even for the computer hard to solve (NP-hard; [7]) and SAT equivalent (**sat**isfiability problem, Sharma et al., 2023). The clever heuristics of cellular decision processes uses decision funnels implemented in signaling cascades to ensure fast decisions. This motivates meta-mathematical considerations that NP problems can never be mapped to P problems (see Supplementary Box 1; only meta-mathematical considerations, no accurate proof).

### Network decision dynamics of platelet activations are well captured by our model

Our observation of the simulation agrees well with all available experimental observations as cited and outlines key nodes and pathways in platelet activation. We could show that upon injury, platelet activation begins with collagen binding to receptors, initiating adhesion via integrin α2β1 and GPVI [17, 18]. This triggers downstream signaling through PLCγ and PI3Kβ, leading to shape change, granule release, and TXA2 generation [18–20]. Activation of PLC increases cytosolic calcium, altering platelet shape and prompting α-granule secretion [21]. Granule release and TxA2 production amplify activation, involving various GPCRs and G proteins [18, 22–25], leading to platelet aggregation through αIIbβ3 activation. This robust response ensures rapid clot formation, regulated by internal and external mechanisms to prevent over-aggregation [26, 27] (More details see supplementary material: box 2 dynamic platelet simulations overview and box 3 enhancing our seven-layered platelet decision model by spatio-temporal modelling with virtual reality using an agent-based model; and supplementary discussion).

### Decision funnels in platelets can shift from a healthy balance to chronic inflammation

We also show how systematic perturbations shift decision-making in platelets, potentially leading to pathophysiological states, a foundation for many diseases, as extensively discussed with literature examples in the supplementary discussion. This feedback loop allows platelets to make rapid, consistent decisions, minimizing errors and maintaining hemostasis. However, under continuous stress, this robustness may prompt thrombo-inflammation due to unceasing activation stimuli, as seen in conditions like diabetes or infection, leading to the loss of endothelial cell (EC) control and hyperactivation of platelets [14, 15]. Our simulations highlight this recurring activation, especially in the absence of EC regulation. Moreover, chronic inflammation alters platelet behavior, switching from thrombosis to promoting inflammation and thrombo-inflammation, elevating the risk of thrombosis [28, 29].

*In silico* analysis of the platelet network shows decision funnels with integration of several pathways. The key example is the preferred Src-kinase pathway for activation, integration of modulatory input and inhition from several pathways. Systems biological analysis shows for them redundancy, error possibilities and error tolearnce, signal augmentation in preferred pathways (signal cascades), but also critical and dispensable nodes, redundant pathway, and shifts of funnels to preferred solutions (free blood flow or thrombosis and hemostasis).

The analysis of decision funnels highlights the critical importance of external feedback in stabilizing decisions. In alignment with SAT theory, the ¾ solution of one cell can be integrated with that of another. When a cell reaches a concurrent conclusion, it receives positive feedback, propelling the decision funnel toward further stabilization and culminating in irreversible aggregation. This process draws a parallel with protein folding, where external factors like water or chaperones similarly influence the funnel.

Moreover, while external conditions are crucial for stabilizing funnels, they can also induce shifts, potentially leading to overstimulation. Likewise, internal dysfunctions, such as those resulting from knockouts (KOs) or mutations, can disrupt the balance between activation and inhibition, causing shifts in the funnel. These shifts may result in either overactivation or a complete lack of activation in platelets, contributing to the onset of various diseases.

### MaxSat analysis demonstrates the heuristics platelets and cells have to apply

In essence, the MaxSat analysis reveals that the potential reasons behind the result of the dynamic analysis lie in the heuristic architecture of the platelet network. This network adeptly simplifies complex (NP-Problems) into more manageable heuristic challenges. It does so by funneling and aggregating information from various pathways to orchestrate a cohesive action (as depicted in Fig. 8).

**Figure 8:**
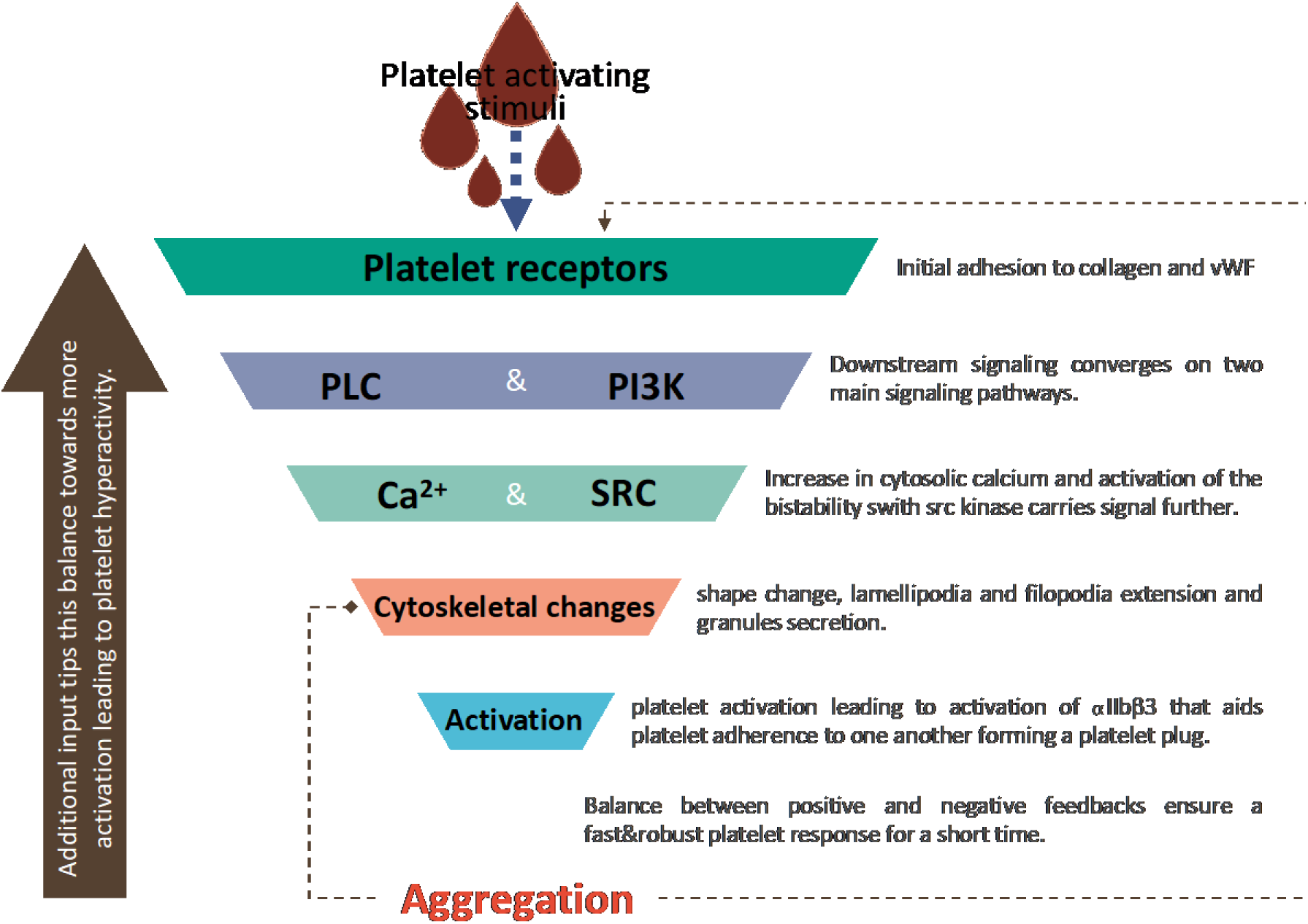
Decision funnel aggregated in several processing layers. Sensors (platelet receptors) measure the environmental change and give environmental feedback. In interconnected downstream cascades, the information is integrated step by step and translated into actions of effectors that induce cytoskeleton changes underlying the process aggregation.

The primary decision funnel is characterized by essential pathways and pivotal nodes, which are crucial components that must invariably be TRUE for the equation (equation 1) to hold. Fig. 8 shows these fundamental pathways along with the critical nodes involved.

This process doesn’t invariably yield the optimal solution but rather provides an approximation, with inherent errors embedded within the decision-making framework (as potentially indicated by errors). Such systems necessitate mechanisms for error correction. In the context of platelets, this is achieved through the incorporation of redundant pathways and the inclusion of dispensable nodes, which refine the decision-making process. Dispensable pathways serve as a safety net for errors in the primary pathway and, alongside dispensable nodes, fine-tune the solution, enhancing the overall outcome beyond the initial ¾ threshold.

Our analysis reveals notable insights by expanding the platelet central network to include feedback loops (refer to Fig. 8, left), which symbolize external environmental feedback, interactions, or updates. Such environmental feedback is often overlooked in protein-protein network simulations. Even with a simplified model incorporated into the network, the influence of feedback on decision funnels is profound. Analogous to protein folding, environmental factors play a crucial role in guiding and stabilizing decision funnels. This can also be viewed as a heuristic approach. While a ¾ solution provides a reasonably effective outcome, direct environmental feedback (either positive or negative) serves to assess and refine the decision, enhancing the precision of subsequent decisions. As a result, the error margin diminishes not by ¼ but by 1/16 when a ¾ solution is combined with another ¾ solution at a subsequent stage. Consequently, rectifying suboptimal decisions becomes more straightforward, as they are promptly adjusted. This dynamic is particularly evident in continuous processes subject to ever-changing environmental conditions. It may lead to oscillatory behavior or environmental scanning, as detailed in [30] or lead to pathologic hyperactivation, for instance in corona infection [31]. These oscillations reflect the system’s constant realignment with environmental stimuli, potentially giving rise to higher-order processes that address complex perceptual challenges [32].

Expanding upon the ¾ solution previously discussed in relation to platelets, our integrated network analysis demonstrates that heuristics and decision funnels provide a robust framework, ensuring clear and consistent decision-making processes [33]. This framework effectively outlines the cell’s primary states, such as the activation and inhibition states observed in platelets. Nonetheless, we have identified decision nodes that exhibit either distinct clarity or ambiguity in their decision-making [33]. Nodes that are decidedly indecisive tend to oscillate frequently between two distinct states—activation or inhibition—characterizing them as bistable. The concept of bistability has been explored, particularly in the context of SRC [4], and is a fundamental characteristic of the RhoA-dependent oscillatory system behavior [30].

### Platelet signalling shows complex cellular decision processes are solved heuristically and motivates to conjecture NP > P

In summary, our sophisticated and innovative combination of analyses demonstrates how cells, exemplified here by platelets, navigate complex challenges. On the one hand, this analytical approach provides a potent tool for unraveling the behavior of various cells and other biological processes. On the other hand, it compellingly argues that living organisms do not solve NP-hard problems by exhaustively searching for perfect solutions. Instead, they employ heuristic strategies that rapidly align with efficient decision-making funnels, further enhanced by mechanisms akin to parallel computing.

This is supported by our theoretical estimation. We propose that NP problems are intrinsically simplified to P problems through the conversion of Boolean SAT problems into linear programming. This suggests that in nature, binary choices are rare, with processes like protein expression and activation occurring over a spectrum of continuous values, allowing for a multitude of satisfactory solutions without necessitating a singular, perfect one (Fig. 9).

**Figure 9:**
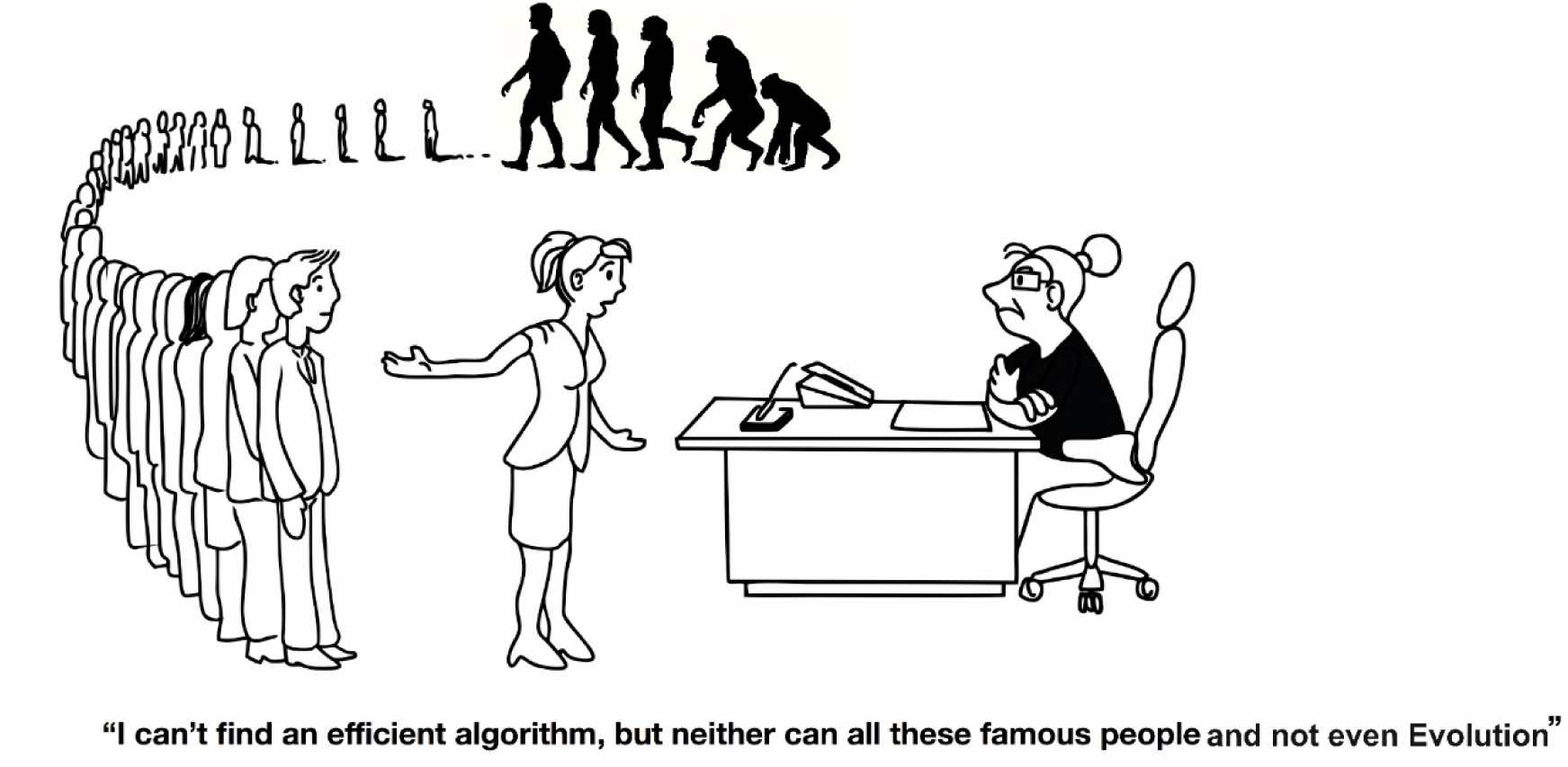
No efficient algorithm mapping NP problems to P problems is known. Illustrated is the evolution of species and its role in finding efficient solutions for solving complex problems. This is a modified version of a famous cartoon by Michael Garey and David S. Johnson in “Computers and Intractability: A Guide to the Theory of NP-Completeness” [9].

We further support this by our conjecture (**Box I**) that posits that no straightforward polynomial-time conversion exists from NP to P problems, owing to the intricate nature of recognizing equivalent solution presentations—a concept rooted in Cantor’s diagonal argument (see metamathematical considerations in Box IS in supplement).

We see that cell biology including platelet signalling doesn’t solve NP problems but rather simplifies them to P problems. Given millions of years of evolution, one could infer that there is no plausible mapping from P to NP possible in the limited decision networks of cells whereas for evolutionary processes and populations even NP problems can be solved using millions of individuals in parallel and finding a close to or even optimal solution in a non-deterministic amount of time. Biology of efficient decision funnels in cells contrasted with the SAT equivalent complexity of cellular decision networks motivates us to believe that NP > P will also be found to be mathematically true. This complex relationship might be further examined using advanced fields like condensed mathematics, then may be proving or clearly rejecting our conjecture which would have significant ramifications for computational complexity theory and the design of decision-making systems.

## Conclusion

We present a seven-layer platelet decision network that aligns well with existing genetic and functional experimental data on platelets. Our model reveals that cellular decision problems are complex (NP-hard and SAT equivalent), but platelet decision cascades are efficient and avoid the stalling seen in NP problems. The network is robust to perturbations, with bistable and multistable nodes playing key roles. Bistable nodes introduce variability to prevent overfitting and getting stuck in local optima, while uncertain decision nodes enhance heuristic solutions by exploring the solution space. This enables optimal and rapid exploration of the complex thrombosis/hemostasis and blood flow dynamics. Platelet decision funnels respond to external stimuli, facilitating swarm intelligence for optimal clot formation and removal, and thrombo-inflammation regulation. The heuristic approach ensures fast decisions (in polynomial time P), avoiding NP problem characteristics. Cellular decision processes hence motivate our meta-mathematical considerations that NP problems can never be mapped to P problems, though we show no mathematical proof for this conjecture.

### Box 1. Some arguments that complex (NP) problems may never be mapped to simple (P) problems

#### Conjecture

- **Hypothesis:** There is no clear polynomial-time (P) computable mapping from NP problems to P problems.
- **Rationale:** The existence of multiple representations for the same solution quality in NP problems (like the Traveling Salesman Problem (TSP)) implies that recognizing equivalent solutions is non-trivial. This is evident even when the solution space is restricted (e.g., travel distances are integers). Thus, identifying a clear P mapping for NP problems appears inherently complex.

#### Supporting Arguments

1. **Complexity of Representation:**

- The essence of the problem lies in the non-obviousness of recognizing identical solutions presented differently.
- Analogous to Cantor’s diagonal argument that establishes a difference in cardinality between the set of real numbers (R) and natural numbers (N), suggesting a fundamental disparity in the scale of complexities between NP and P.
2. **Condensed Mathematics as a Tool:**

- The field of condensed mathematics offers methods to identify and equate closely related representations (e.g., recognizing 0.999… as 1).
- This methodology might be applicable to computational problems like SAT, potentially offering a new approach to understanding the relationship between NP and P by efficiently recognizing equivalent solution representations.
3. **Meta-mathematical Considerations** (see Suppl. Box 1):

- Beyond mathematical formalism, philosophical and meta-mathematical considerations, such as the implications of the halting problem, suggest a higher intrinsic complexity in NP problems compared to P problems.
4. **Practical Observation in Biological Systems:**

- Despite the theoretical complexity, living cells demonstrate a preference for efficient heuristics over precise solutions to NP problems, reflecting a practical approach to decision-making in complex, real-world systems.

#### Mathematical Formulation

While a rigorous mathematical proof of this conjecture is beyond the scope of this response, the conjecture can be formalized within the framework of computational complexity theory. The challenge lies in formally demonstrating that:

- No efficient (polynomial-time) algorithm can transform every problem instance in NP into an equivalent instance in P such that the solutions correspond and are computationally feasible to identify.
- The inherent complexity in recognizing equivalent solutions (even under constrained conditions) contributes to the non-existence of a clear polynomial-time reduction from NP to P.

This formulation sets the stage for a profound investigation in computational complexity theory, potentially involving advanced mathematical tools and concepts from fields like condensed mathematics, and bearing significant implications for both theoretical computer science and practical problem-solving in various domains, including biology.

## Abbreviations

NP: non-deterministic complete
P: always polynomial time taking
SAT: Boolean satisfiability problem.

## Acknowledgements

We thank Jan Grundheber, Tobias Schmidt and Dennis Eckert for sharing their platelet videos (virtual reality, agent-based simulation) with us.

The authors gratefully acknowledge the support by Deutsche Forschungsgemeinschaft (DFG) Project number 3 74031971 – TRR 240 [HS: A3][ÖO, JB, TD, KH:INF] platelet start model. Project number DFG project number 492620490 - SFB 1583/INF (TD, JB: Decision steps, refined platelet model, inflammation). TD/JP acknowledge in addition project number 210879364 – TRR 124 [B1/B2] (game theory, conjecture/ SAT equivalence).

## Author contributions

JP, JB, ÖO did all bioinformatical analysis. JP did work out the SAT proof and transferred it to the platelet protein network. TD worked on the meta-mathematics and the conjecture. JB, MA and ÖO constructed the platelet signaling network according to the experimental data and simulated. TD lead and guided the study.

JP, JB, ÖO, MK, MS, KGH, HS and TD were all involved in the data analysis. MK, MS, KGH, SvM, HS provided expert analysis on the network behavior.

JP, JB, ÖO and TD were involved in drafting the paper which was followed by editing and comments by all authors. All authors read and approved the final manuscript.

Formal contributions (in bold selected):

JP: concept, curation, formal, funding, invest, method, admin, soft, super, vali, visu, ori, edit JB: concept, curation, formal, funding, invest, method, admin, soft, super, vali, visu, ori, edit ÖO: concept, curation, formal, funding, invest, method, admin, soft, super, vali, visu, ori, edit MK: concept, curation, formal, funding, invest, method, admin, soft, super, vali, visu, ori, edit MS: concept, curation, formal, funding, invest, method, admin, soft, super, vali, visu, ori, edit MA: concept, curation, formal, funding, invest, method, admin, soft, super, vali, visu, ori, edit SvM: concept, curation, formal, funding, invest, method, admin, soft, super, vali, visu, ori, edit KGH: concept, curation, formal, funding, invest, method, admin, soft, super, vali, visu, ori, edit HS: concept, curation, formal, funding, invest, method, admin, soft, super, vali, visu, ori, edit TD: concept, curation, formal, funding, invest, method, admin, soft, super, vali, visu, ori, edit

## Availability of data and materials

All results are contained in the manuscript and its supplementary files including all details on the modeling involved and all experimental data for the study.

## Supporting information

**Supplementary File 1:** Supplementary Tables S1-2, Supplementary Figures S1-10, Supplementary Introduction, Supplementary Results, Supplementary Discussion.

**Supplementary File 2:** Platelet central cascade model with feedback and env. inputs in graphml format.

**Supplementary File 3:** Platelet central cascade model without feedback and env. inputs in graphml format.

**Supplementary File 4:** Platelet central cascade model with diseases in graphml format.

### Supplementary videos

i. A video using Blender on platelet vessel flow (video S1.mp4).
ii. A video using Unity on platelet activation (video S2.mkv).
iii. A video of what happens inside a cell while interacting with the environment using Jimena (video S3.mp4).
iv. The activation of the inhibitory cyclic nucleotide pathway and platelet behavior change such as shape changes (video S4.mp4).

