## Supplemental Materials for "Thrombo-inflammation analyzed in a validated seven-layer platelet decision model: cellular decisions are tough problems fast and heuristically solved"

### Contents

Analyze platelet signaling with a SAT solver ................................................................... 2
The SAT problem ............................................................................................................... 2
Platelet signaling cascade translated into a Boolean function or Platelet signaling cascade and model applied to show that cellular decisions represent an SAT problem formally. …….. 2
Integer program formulation of MaxSat (IP) ..................................................................... 4
Computational Analysis of Biological Networks using SAT and MAX-SAT Solvers ....... 5
Computational scripts used……………………………………6
Supplementary Introduction ............................................................................................... 6
Comparing tough NP problems to more reliable solvable P problems (p.6)
**Supplementary Results** ..................................................................................................... 7
Supplementary Box I: Meta-mathematical considerations comparing NP and P problems.................................................................................................................................... 7
Supplementary Box 2: Simplified Overview of Dynamic Platelet Simulation Results ...... 9
Supplementary Box 3: Enhancing our seven-layered platelet decision model by virtual reality and agent-based models....................................... ……………………………………10
Supplementary Table S1 (p. 13), Table S2 (p.15), Table S3 (p. 31).
Supplementary Figures .......................................................................................................15
Supplementary Discussion...................................................................................................22
Supplementary References (p. 31)

### Analyze platelet signaling with a SAT solver

#### The SAT problem

SAT is a decision problem that decides whether a Boolean formula, written in the conjunction normal form (CNF), can be evaluated as true. Let $x_i$ be a Boolean variable, which takes only 0 (false) or 1 (true) values. Let ${l_i}^j$ be the literal associated with the Boolean variable $x_i$ in the clause $j$. ${l_i}^n$ its either $x_i$, $\neg x_i$ or 0. The literal either takes the value of the Boolean variable or its negation or 0 (false). This last option represents that the Boolean variable is not part of a clause. A clause in the CNF form is a disjunction of literals. A complete Boolean expression is a conjunction of clauses, as seen in **equation S1**.

$$f(\boldsymbol{x}) = \left({l_1}^1 \vee {l_2}^1 \vee \dots \vee {l_n}^1\right) \wedge \left({l_1}^2 \vee {l_2}^2 \vee \dots \vee {l_n}^2\right) \wedge \dots \wedge \left({l_1}^j \vee {l_2}^j \vee \dots \vee {l_n}^j\right) \wedge \dots \qquad (eq.1)$$

**equation S1**. The general form of a Boolean expression in CNF form.

Accordingly, SAT decides whether there exists a set $\boldsymbol{x}$ of Boolean variables that satisfies (evaluates to true) such an expression f(x). Usually, we set the maximum number of variables allowed in each part of a SAT problem. Because of this, we often call the SAT problem the K-SAT problem. In this name, 'K' represents the largest number of variables any part can have. The 2-SAT problem is a P problem, but 3-SAT and all other bigger forms are NP-complete.
Another problem is the Max-SAT problem. It is defined by whether the Boolean expression is satisfiable or not. The idea is to find the combination of Boolean variables $\boldsymbol{x}$ such that the maximum number of clauses possible is satisfied. These problems are also NP-complete.

#### Platelet signaling cascade translated into a Boolean function or Platelet signaling cascade and model applied to show that cellular decisions represent an SAT problem formally.

To interpret this signaling cascade as an SAT problem, we first need to transform it into a Boolean network. This transformation is straightforward simply by assuming that each protein in the network is a Boolean variable and it has only two possible states: 0 (inactive/false) or 1 (active/true).
Now, the edges connecting proteins mean that the protein on the ending side depends on the protein on the starting side. This means that it can be activated or inhibited by the parent protein of the edge depending on the shape of the edge termination. An arrow-shaped edge is used for activation, and an orthogonal short line is used for inhibition. For example, STIM1 activates ORAI1, which in Boolean terms would be written as $STIM1 = ORAI1$. As an inhibitory example, see that P2Y12 inhibits AC. In this case, the Boolean expression would be $AC = \neg P2Y12$.

If multiple edges arrive at a protein, it means it is controlled by these respective incoming signals. Overall, this yields a logical OR among all the parent proteins. For example, the Boolean formula for the IP3R node in **Fig. 2** is:

$IP3R = (IP3 \lor \neg PKA \lor \neg PKG)$.

Here, the usage of OR is sufficient for the modeling (Karl and Dandekar, 2015a): In cellular signaling, it is common to encounter redundant activation pathways. Essentially, this operates much like an 'OR' gate, where multiple routes can independently lead to the activation of the same function or response. Moreover, the reaction of multiple molecules at once is rare. Usually, compounds are formed in a sequential form. If one wants to represent an AND, the usual trick is to create an artificial intermediate inhibitory node. As a result, the PPI network can be transcribed into a Boolean expression using this simple convention.

As outlined above, several processes must be initiated to have a successful clot formation process that prevents the organism from bleeding. We included 5 of those processes in the network: Granule release (GR), Aggregation (Ag), Stress fiber formation (SFF), Lamellipodia (La), and Shape change (SC). We refer to the coagulation process as CP. The full CP requires each of the five processes. The Boolean expression corresponding to that situation would be as follows:

$$CP = GR \land Ag \land SFF \land La \land SC$$

We can further expand this Boolean expression by replacing each of the processes involved with the corresponding set of activating parent nodes. In that case, the coagulation process would be described as follows:

$$CP = (RAC1) \land (a2b3) \land (Ca2^{+} \lor ROCK1) \land (RAC1) \land (SFF \lor La)$$

Following the procedure of replacing the nodes with their Boolean activation equation, we would obtain:

$$CP = (RAP1) \land (TALIN1) \land (P2X1 \lor ORAI1 \lor Mg2^{+}{}_{trans} \lor IP3R \lor TRPC6 \lor \neg Mg2^{+} \lor RHOA) \land (RAP1) \land (Ca2^{+} \lor ROCK1 \lor RAC1)$$

Then, recursively following the same procedure, we obtain:

$$CP = (PKC \lor CALDAG \lor \neg PKA \lor \neg PKG) \land (RAP1) \land (ATP \lor STIM1 \lor PKC \lor IP3 \lor \neg PKA \lor \neg PKG \lor DAG \lor \neg PKC \lor \neg(TRPM7 \lor Mg2^{+}{}_{trans}) \lor ARHGEF1) \land (PKC \lor CALDAG \lor \neg PKG \lor \neg PKA) \land (P2X1 \lor ORAI1 \lor Mg2^{+}{}_{trans} \lor IP3R \lor TRPC6 \lor \neg Mg2^{+} \lor RHOA \lor RAP1)$$

Finally, by solving the negated OR, we get:

$$CP = (PKC \lor CALDAG \lor \neg PKA \lor \neg PKG) \land (RAP1) \land (ATP \lor STIM1 \lor PKC \lor IP3 \lor \neg PKA \lor \neg PKG \lor DAG \lor \neg PKC \lor \neg TRPM7 \lor ARHGEF1) \land (\neg {Mg2^{+}}_{trans}) \land (PKC \lor CALDAG \lor \neg PKG \lor \neg PKA) \land (P2X1 \lor ORAI1 \lor {Mg2^{+}}_{trans} \lor IP3R \lor TRPC6 \lor \neg Mg2^{+} \lor RHOA \lor RAP1)$$

Notice that this expression is not final and that we could go on replacing different nodes with their more complex, respective activating function, but there is no need to go further. The expression we observe is already a clear example of a Boolean expression in the CNF form.

Observing this expression allows us to formulate the platelet signaling cascade problem as an NP problem, precisely the MaxSAT problem. The idea is that to perform the coagulation process, the organism needs to activate each of the processes involved (e.g. aggregation, stress fiber formation, adhesion). It does not necessarily need to activate all of them simultaneously, but it should activate as many of them as possible since bleeding can be deathly very quickly. Hence, we assume that the organism is trying to find the combination of active and nonactive proteins which achieves the maximum number of active coagulation processes. In the best-case scenario, all of them are active at once. This is the exact definition of the MaxSAT problem, as explained above. Accordingly, we are trying to find the Boolean variables (proteins) combination that maximizes the number of satisfied clauses (coagulation processes).

In the same way, we proceeded with the platelet signaling cascade, it is possible to interpret any PPI network as a MaxSAT NP problem. Nevertheless, as stated before, our organism deals with such issues at every moment and seems to solve them accurately and quickly. We will now try to understand how this happens and what we can learn from it.

#### Integer program formulation of MaxSat (IP)

$$\max \sum_{C_j \in \mathfrak{C}} w_j z_j$$

$$subject\ to:$$

$$\sum_{i \in I_j^{+}} y_i + \sum_{i \in I_j^{-}} (1 - y_i) \geq z_j \qquad \forall\, C_j \in \mathfrak{C}$$

$$y_i \in \{0,1\} \qquad \forall\, i \in \{0,1,\dots,n\}$$

$$z_j \in \{0,1\} \qquad \forall\, C_j \in \mathfrak{C}$$

$C_j$ is the clause $j$ in the set of all clauses $\mathfrak{C}$. Where, $w_j$ is the weight assigned to clause $j$. $z_j$ is 1 if the clause $j$ is satisfied and 0 otherwise. $y_i$ is a literal in the clause which equals 1 if the variable $x_i$ is true and 0 if $x_i$ is false. The objective function is to maximize the value obtained from satisfying clauses with their respective assigned weight. If we set all weights to 1, the program tries to satisfy as many clauses as possible. The first constraint ensures that at least one variable is positive or a negated variable is negative because the clauses are conjunctions of variables, so one *true* value is enough to satisfy the clause. The second restriction assures the Boolean nature of the variables. The third condition ensures that the clauses will be evaluated as *true* or *false.*

#### Computational Analysis of Biological Networks using SAT and MAX-SAT Solvers

In this study, we employed SAT and MAX-SAT solvers to model and analyze the logical structure of the platelet network. Specifically:

- **SAT Solvers:** These were used to determine the exact conditions under which the platelet network is activated. This involves identifying scenarios where the Boolean formula representing the network is fully satisfied.
- **MAX-SAT Solvers:** These solvers provided insights into the network's behavior when not all conditions can be simultaneously met. They helped highlight the most critical components and pathways by identifying the maximum number of clauses that can be satisfied concurrently.

Heuristic and Decision Funnel Analysis

We conducted a comparative analysis of the solutions obtained from SAT and MAX-SAT solvers to:

- Identify the heuristics and decision funnels employed by the platelet network.
- Understand the network's preferred pathways, critical nodes, and potential error-prone routes, offering insights into the network's decision-making processes under various physiological and pathophysiological conditions.

Applications in Systems Biology

Our methodology was applied in several key areas within systems biology:

- **Critical Node Identification:** We pinpointed essential components within the platelet network, assessing the impact of the presence or absence of these components on the overall functionality of the network.
- **Pathway Redundancy and Robustness:** Our approach helped reveal redundant pathways and their contributions to the network's robustness, enhancing our understanding of how biological systems maintain functionality under stress or perturbations.
- **Error Analysis and Pathophysiological Insights:** We identified error-prone pathways and preferred states within the network, providing crucial insights into potential

pathophysiological mechanisms. This analysis is instrumental in guiding future experimental and therapeutic strategies.

**Computational script used**

Here is the python script for the SAT solver. It is purposely constructed to have a simple code, simply the definition of the problem and the call to the solver.

```
from pysat.solvers import Glucose3

#CP=(PKC ∨CALDAG ∨ ¬ PKA∨ ¬ PKG)  ∧
#  (RAP1) ⋀ (ATP ⋁ STIM1 ⋁ PKC ∨ IP3∨DAG ∨ ¬PKC ∨ ¬TRPM7 ∨ ARHGEF1 )  ∧( 〖¬
 〖Mg2〗 ^+〗 _trans)  ⋀ (PKC ∨CALDAG∨¬PKG∨¬PKA) ⋀   (P2X1 ⋁ ORAI1 ⋁  〖 〖Mg2
〗 ^+〗 _trans  ∨ IP3R ∨TRPC6∨
# ¬ 〖Mg2〗 ^+  ∨RHOA ∨ RAP1)
g = Glucose3()

g.add_clause([1,2,-3,-4])
g.add_clause([5])
g.add_clause([6,7,1,9,-3,-4,12,-1,-10,8])
g.add_clause([15])
g.add_clause([1,2,-4,-3,])
g.add_clause([13,14,15,11,16,-17,18,5])


print(g.solve())
print(g.get_model())
```

### Supplements to the Introduction

#### Comparing tough NP problems to more reliable solvable P problems

An NP (Non-deterministic Polynomial time) problem is defined in computational complexity theory as a problem that cannot be solved in polynomial time by a deterministic machine, but one can check in polynomial time whether a given solution is valid. As opposed to NP problems, there are P (Polynomial time) problems that a deterministic machine can solve in polynomial time. As a result of this, the mathematical term deterministic machine refers to a normal computer, and the expression "to solve in polynomial time" refers to a number of calculations a computer must make to solve a problem that grows polynomially with the size of the problem as opposed to, for example, linearly or exponentially.

NP problems are impactful, and people often refer to them as unsolvable. In theory, NP problems are solvable, but it may take an unpractical amount of time to find a solution, even for relatively small instances of the problem. A common example is the traveling salesperson problem (TSP). A vendor must find the shortest path to visit a set of cities. When attempting to solve the problems with a brute force strategy by testing all possible combinations, the number of calculation steps grows with the factorial number of cities ($O(n!)$). Consequently, even for a small example of only 15 cities, one would have to make over 1.3 trillion calculation steps. This is clearly not feasible, and the calculation problem worsens as the number of cities increases.

One should also mention that an NP-complete class is within the set of NP problems. These are problems that are believed to be in NP and mapped into one another, meaning that if someone found a polynomial time solution for one of them, it would imply that all of those would be solvable in polynomial time.

#### Supplements to the Results

##### Supplementary Box I: Meta-mathematical considerations comparing NP and P problems

Understanding the relationship between complex NP problems and simpler P problems is a big challenge in mathematics. Here, we're sharing some high-level thoughts on how we might begin to understand this relationship. These ideas are inspired by our findings in biology and are meant to be a starting point, not a detailed proof.

**Idea of the Proof:** Our studies in biology show that cells don't solve extremely complex problems directly. Instead, they deal with simpler problems efficiently. This observation leads us to think:

**Part I: Comparing the range of solutions for NP and P Problems**

- It seems that NP problems have a wider range of possible solutions compared to P problems, making it hard to find a direct way to convert NP problems into P problems.

**How do we explore this?** We draw parallels with a similar problem in set theory, where we compare different types of number sets, like the natural numbers (N), whole numbers (Z), rational numbers (Q) which are all infinitely large but in a manageable way, and compare them to the real numbers (R) which represent a much larger kind of infinite.

**Similar Problem: Comparing Different Sets of Numbers**

- It's easy to show that sets N, Z, and Q have a similar size; you can match their elements one-to-one.
- However, set R is a different type of infinite, significantly larger.

**Proof:** We reference Cantor's diagonal argument, a famous proof (see, e.g., "The Discovery of the Infinite" by David Foster Wallace (Wallace, 2009).

**But there's a counterargument:**

- Some mathematicians argue against Cantor's method, saying that infinite series like this don't really exist. They believe that only numbers or sequences that we can fully describe should count. If we follow this line of thinking, set R would be considered similar in size to sets N, Z, and Q.

**Part II: Applying a meta-mathematical argument of analogy to our original problem (P vs NP)**

- You might think about creating a complete list of solutions for an NP problem, effectively turning it into a P problem. This approach works as long as you don't need infinitely long solutions or sequences of steps. This is the viewpoint of constructivism.

**But a major challenge is the "halting problem":**

- With some problems (especially in NP), you can't know in advance if the problem will ever be solved.
- So, the question is: "Can the solutions for NP problems be neatly organized into P problems?" Given the nature of NP problems and the halting problem, constructivism doesn't seem to fit well here.

**Considerations**: Suppose we would have a mapping of all NP problems to P problems, showing that NP = P analogous to a countable set. Cantor´s diagonal proof would then take one element of each entry of the list (imagine the list of city itineraries in TSP) and show that this journey is not in the list of cardinality P and could add like this ever more journeys, showing that the journey list is non-countable long. However, this is only allowed if also very long or may be even never ending journey representations are allowed. In constructivism, this would be forbidden. In NP problems, this is allowed ("halting problem"). Hence, this strongly suggests the cardinality of NP problems is clearly larger then P due to ever more only *may be similar* long journey representations can easily be created. However, this is an argument of analogy, a formal proof is not given here.

**Conclusion:** Our thoughts, combined with what we know from biology, suggest that NP problems are inherently more complex, and you can't simplify them into P problems easily. This indicates that there's no straightforward way to map every NP problem to a P problem, making NP problems fundamentally larger or more complex than P problems.

#### Supplementary Box 2: Overview of Dynamic Platelet Simulation Results

This section reviews our computer simulations to understand how platelets behave under different conditions.

**Simulation Tool & Model:**

- We used the software Jimena and a specific model structure (platelet_network.graphml in supplementary material) to run our simulations. This helped us see how platelets make decisions and interact with each other in their signaling processes.

**Key Findings and Simulations:**

First of all, the model captures healthy state, activation and inhibition as modelled and published previously (Mischnik et al., 2014; Breitenbach et al., 2022). However, we took now into account also details of pathophysiological states and how the decision network would respond. This revealed the 7 decision layers and decision funnels helping fast reactions. Regarding pathophysiology, we looked at:

1. **Damage Response:**
   - We observed how damage to the epithelial cells (cells lining blood vessels) leads to the exposure of collagen (a protein), which then triggers platelets to activate and respond to the injury.
   - We looked at various situations:
     - Constant damage to the epithelial cells.
     - Complete absence of the epithelial cell layer.
     - Effects of chronic inflammation on platelets.
     - Impact of atherosclerosis (artery hardening due to plaque) on platelets, both with and without cell damage.
     - Influence of inflammatory arthritis on platelets, comparing situations with and without cell damage.
     - How aging affects platelets in the above conditions.

- These situations were simulated and visualized in different figures (referred to as Figure 1, S4, S5, S6, S7, S8, and S9).

2. **Impaired Signaling:**
   - We also studied what happens when platelets can't signal properly. For example:
     - When TRPM7 (a protein involved in magnesium and calcium signaling) is absent.
     - When platelets are isolated and don't receive external signals.
   - These conditions were also visualized in figures (Figure 1, S1, S2, S3).
3. **Genetic Differences:**
   - We simulated how platelets with different genetic makeup respond to damage or other stimuli compared to normal (wild-type) platelets. This included:
     - Platelets deficient in GPVI, a protein involved in their activation.
     - Platelets from individuals with Bernard-Soulier Syndrome, which affects how platelets stick together.
     - Platelets from individuals with Glanzmann Thrombasthenia, another condition affecting platelet function.
   - These scenarios are illustrated in figures (Figure S10, S11, S12).

**Conclusion:**

- Our model captures how platelets behave, starting from healthy state and then taking into account various factors modifying signaling responses like genetic differences, absence of certain proteins, and different disease conditions.
- We found that both the central activation process of platelets and the additional inputs (like genetic factors or disease states) are crucial to understanding how platelets respond to their environment. A decision funnel arodun the central src kinase orchestrates rapid activation, modifications include inhibitory input via cyclic nucleotides and modulation via calcium and magnesium signaling.
- Our *in silico* model mimicks the heuristic, fast and clear decision-making of platelets by including all signaling proteins and components known to have an impact

according to available genetic, functional or pathophysiological data. This avoids the complexity of the tough (NP) problems in our simulations. Also, we noted that in normal (wild-type) platelets, the signaling process doesn't get stuck or halts unexpectedly (neither in experiments no in the simulations).

**Supplementary Box 3: Enhancing our seven-layered platelet decision model by virtual reality and agent-based models**

Virtual reality (VR) allows to enhance our seven-layered platelet decision model: The seven-layered signaling network is translated in its key properties into an agent-based model (ABM) where one agent represents one platelet. Using the VR we can simulate the behavior of many platelets and how they move in space and time. The VR allows hence to model and assess respective influences of the platelets, the surrounding tissue and arising pathophysiological effects. In general, a VR environment using ABM is particularly strong to elucidate spatio-temporal sequences and events. To be as realistic as possible, we integrate also image information, for example the texture of different objects, e.g. platelets, endothelial cells, vasculature (which then, in the next refinement steps influence platelet behavior). A couple of videos illustrate this:
(i) A video by Jan Grundheber using Blender on platelet vessel flow (video S1.mp4).
(ii) A video by Tobias Schmidt and Dennis Eckert using Unity on platelet activation (video S2.mkv).
(iii) A video by Johannes Balkenhol of what happens inside the platelet cell while interacting with the environment using Jimena (video S3.mp4).

For the dynamic simulations of the seven-layered decision network we use our dynamic network analysis software, Jimena. To become more realistic and allow a VR this is being translated from Java into C# to enhance its functionality in Godot and other game engines to establish a VR. This translation is a critical step in integrating sophisticated signaling networks into our virtual environment, specifically tailored to the study of platelets. By this we translate the validated signaling networks of the platelet into our virtual models looking at many platelets and their behavior in space and time. We next create machine-readable network formats for advanced analysis.

*Integration of Networks into Godot* (preferred to Unity and Blender as Godot is open source; but the procedures are similar): The networks, delivered in GraphML format, are seamlessly integrated into Godot using a specially designed C# language plugin, incorporating the Jimena Software GraphML importer. The core of our network simulation is the SQUAD method, which intuitively translates network topology into dynamic networks, altering activation patterns over time.

*Simulations for Network Control and Behavior of many platelets*: We have already developed detailed Boolean network models of central platelet signaling, inhibition and modulation. The platelet receptors and secreted platelet factors modelled, allow platelets to influence each other.

To give two central examples we currently are studying: using the Godot engine via Java and C# we turn our 7-layered model into an ABM and to reproduce the cAMP effect in the platelets in VR, and also the simple effect that the inflammatory cytokines increase in the tissue after the thrombus in the brain.

In particular, the spatio-temporal model of the ABM allows to study in a detailed way thrombus formation and packing of the platelets to and within the thrombus. cAMP concentrations are comparatively high (inhibitory for platelet activation; see Figure 2 in paper and Breitenbach et al., 2022) in the outside regions of the thrombus, but low (and hence the thrombus is more solid inside the thrombus). Critical protein interactions and concentrations, for instance the activation of the inhibitory cyclic nucleotide pathway is incorporated by Godot match and case statements.

```
var cAMP: int = 1
match x:
	1:
		# If cAMP == 1, run this block of code
	_:
		 # If nothing matches, run this block of code
```

The activation of the inhibitory cyclic nucleotide pathway are simulated and translated into receptor activation which drive via the network logic (e.g. in Fig. 2) the behavior change such as shape changes as demonstrated in video S4. This video is no *in silico* simulation but original validation data from an experiment: A time-lapse video of intravital confocal microscopy of the superior sagittal sinus, staining of vasculature (violet, BSA-A546 and anti-CD105-A546) and platelets (green, anti-GPIV A488), showing platelets during vessel thrombosis in the brain.

The ABM allows to monitor the thrombus formation and how antithrombotic or prothrombotic factors influence this including surface effects on thrombus and endothel and effects of the endothelial cells.

Similarly, the spatio-temporal model of the ABM allows to study in a detailed way how inflammatory changes in the brain tissue happen in stroke: We can collect from the model detailed insights into cytokine effects before and after the thrombus on the brain tissue. In particular we can study the extent of the high inflammation penumbra in stroke and how this can be modulated (e.g. by drugs and T-cells) and compare tissue before and after the occlusion or inside and outside the stroke region.

With both examples, we can also show very clearly what we gain compared to our single platelet decision model (single model, fully discussed in this paper): With the spatiotemporal ABM model, we can emphasize the communication between the platelet agents as a further emergent effect, which can then be investigated in the VR model during thrombus formation, but never in our Boolean model as well as the inflammatory stimulation of the tissue by the platelets during stroke (modelled by agents representing endothelial cells).

### Supplementary Tables

**Table S1:** Details of simulations, perturbations for each simulation are given for each figure.

| Simulation | Perturbation | Figure | Comment |
|---|---|---|---|
| Quiescent state simulation | damage/injury: Value 0 from 0 to 1000 | Figure 3 | WT |
| Damage to the EC layer | damage/injury: Value 1 from 0 to 5 | Figure 4 | WT |
| Threshold simulation in vivo | damage/injury: Values 1 from 0 to 5<br>TRPM7: Value 0.5 from 0 to 1000 | Figure 5A | WT |
| | damage/injury: Value 0.4 from 0 to 5<br>TRPM7: Value 0.5 from 0 to 1000 | Figure 5B | WT |
| | damage/injury: Value 0.3 from 0 to 5<br>TRPM7: Value 0.5 from 0 to 1000 | Figure 5C | WT |
| | damage/injury: Value 0.2 from 0 to 5<br>TRPM7: Value 0.5 from 0 to 1000 | Figure 5D | WT |
| | damage/injury: Value 0.1 from 0 to 5<br>TRPM7: Value 0.5 from 0 to 1000 | Figure 5E | WT |
| | damage/injury: Values 1 from 0 to 5<br>TRPM7: Value 0 from 0 to 1000 | Figure 5F | KO |
| | damage/injury: Value 0.4 from 0 to 5<br>TRPM7: Value 0 from 0 to 1000 | Figure 5G | KO |
| | damage/injury: Value 0.3 from 0 to 5<br>TRPM7: Value 0 from 0 to 1000 | Figure 5H | KO |
| | damage/injury: Value 0.2 from 0 to 5<br>TRPM7: Value 0.5 from 0 to 1000 | Figure 5I | KO |
| | damage/injury: Value 0.1 from 0 to 5<br>TRPM7: Value 0 from 0 to 1000 | Figure 5J | KO |
| Threshold simulation in vitro | damage/injury: Values 1 from 0 to 5<br>TRPM7: Value 0.5 from 0 to 1000 | Figure 6A | WT |
| | damage/injury: Value 0.4 from 0 to 5<br>TRPM7: Value 0.5 from 0 to 1000 | Figure 6B | WT |
| | damage/injury: Value 0.31 from 0 to 5<br>TRPM7: Value 0.5 from 0 to 1000 | Figure 6C | WT |
| | damage/injury: Value 0.2 from 0 to 5<br>TRPM7: Value 0.5 from 0 to 1000 | Figure 6D | WT |
| | damage/injury: Value 0.11 from 0 to 5<br>TRPM7: Value 0.5 from 0 to 1000 | Figure 6E | WT |
| | damage/injury: Values 1 from 0 to 5<br>TRPM7: Value 0 from 0 to 1000 | Figure 6F | KO |
| | damage/injury: Value 0.4 from 0 to 5<br>TRPM7: Value 0 from 0 to 1000 | Figure 6G | KO |
| | damage/injury: Value 0.31 from 0 to 5<br>TRPM7: Value 0 from 0 to 1000 | Figure 6H | KO |
| | damage/injury: Value 0.2 from 0 to 5<br>TRPM7: Value 0.5 from 0 to 1000 | Figure 6I | KO |
| | damage/injury: Value 0.11 from 0 to 5<br>TRPM7: Value 0 from 0 to 1000 | Figure 6J | KO |
| Atherosclerosis | kinase: Value 0.5 from 0 to 1000<br>atherosclerosis: Value 0 from 0 to 1000 | Figure 7A | WT |
| | kinase: Value 0.5 from 0 to 1000<br>atherosclerosis: Value 0 from 0 to 1000<br>damage/injury: Value 1 from 0 to 5 | Figure 7B | WT |
| | kinase: Value 0.5 from 0 to 1000<br>atherosclerosis: Value 0.4 from 0 to 1000 | Figure 7C | WT |
| | kinase: Value 0.5 from 0 to 1000<br>atherosclerosis: Value 0.4 from 0 to 1000<br>damage/injury: Value 1 from 0 to 5 | Figure 7D | WT |
| | kinase: Value 0.5 from 0 to 1000<br>atherosclerosis: Value 0.52 from 0 to 1000 | Figure 7E | WT |
| | kinase: Value 0.5 from 0 to 1000<br>atherosclerosis: Value 0.52 from 0 to 1000<br>damage/injury: Value 1 from 0 to 5 | Figure 7F | WT |

| | | | |
|---|---|---|---|
| | kinase: Value 0.5 from 0 to 1000<br>atherosclerosis: Value 0.7 from 0 to 1000 | Figure 7G | WT |
| | kinase: Value 0.5 from 0 to 1000<br>atherosclerosis: Value 0.7 from 0 to 1000<br>damage/injury: Value 1 from 0 to 5 | Figure 7H | WT |
| Inflammatory arthritis (IA) | kinase: Value 0.5 from 0 to 1000<br>IA: Value 0.5 from 0 to 1000 | Figure S6A | WT |
| | kinase: Value 0.5 from 0 to 1000<br>IA: Value 0.5 from 0 to 1000<br>damage/injury: Value 1 from 0 to 5 | Figure S6B | WT |
| | kinase: Value 0.5 from 0 to 1000<br>IA: Value 0.52 from 0 to 1000 | Figure S6C | WT |
| | kinase: Value 0.5 from 0 to 1000<br>IA: Value 0.52 from 0 to 1000<br>damage/injury: Value 1 from 0 to 5 | Figure S6D | WT |
| | kinase: Value 0.5 from 0 to 1000<br>IA: Value 0.58 from 0 to 1000 | Figure S6E | WT |
| | kinase: Value 0.5 from 0 to 1000<br>IA: Value 0.58 from 0 to 1000<br>damage/injury: Value 1 from 0 to 5 | Figure S6F | WT |
| Aging | kinase: Value 0.5 from 0 to 1000<br>aging: Value 0.5 from 0 to 1000 | Figure S7A | WT |
| | kinase: Value 0.5 from 0 to 1000<br>aging: Value 0.5 from 0 to 1000<br>damage/injury: Value 1 from 0 to 5 | Figure S7B | WT |
| | kinase: Value 0.5 from 0 to 1000<br>aging: Value 0.52 from 0 to 1000 | Figure S7C | WT |
| | kinase: Value 0.5 from 0 to 1000<br>aging: Value 0.52 from 0 to 1000<br>damage/injury: Value 1 from 0 to 5 | Figure S7D | WT |
| | kinase: Value 0.5 from 0 to 1000<br>aging: Value 0.58 from 0 to 1000 | Figure S7E | WT |
| | kinase: Value 0.5 from 0 to 1000<br>aging: Value 0.58 from 0 to 1000<br>damage/injury: Value 1 from 0 to 5 | Figure S7F | WT |
| GPVI-deficiency | kinase: Value 0.5 from 0 to 1000<br>damage/injury: Value 1 from 0 to 5 | Figure S8A | WT |
| | kinase: Value 0.5 from 0 to 1000<br>damage/injury: Value 1 from 0 to 5<br>GPVI: Value 0 from 0 to 1000 | Figure S8B | GPVI-deficiency |
| Bernard-Soulier Syndrome (BSS) | kinase: Value 0.5 from 0 to 1000<br>vWF: Value 1 from 0 to 5 | Figure S9A | WT - vWF stimulation |
| | kinase: Value 0.5 from 0 to 1000<br>vWF: Value 1 from 0 to 5<br>GP1b-IX-V: Value 0 from 0 to 1000 | Figure S9B | BSS - vWF stimulation |
| | kinase: Value 0.5 from 0 to 1000<br>collagen: Value 1 from 0 to 5 | Figure S9C | WT - collagen stimulation |
| | kinase: Value 0.5 from 0 to 1000<br>collagen: Value 1 from 0 to 5<br>GP1b-IX-V: Value 0 from 0 to 1000 | Figure S9D | BSS - collagen stimulation |
| | kinase: Value 0.5 from 0 to 1000<br>ADP: Value 1 from 0 to 5 | Figure S9E | WT - ADP stimulation |
| | kinase: Value 0.5 from 0 to 1000<br>ADP: Value 1 from 0 to 5<br>GP1b-IX-V: Value 0 from 0 to 1000 | Figure S9F | BSS - ADP stimulation |
| Glanzmann Thrombasthenia (GT) | kinase: Value 0.5 from 0 to 1000<br>damage/injury: Value 1 from 0 to 5 | Figure S10A | WT |
| | kinase: Value 0.5 from 0 to 1000<br>damage/injury: Value 1 from 0 to 5<br>$\alpha IIb\beta 3$: Value 0 from 0 to 1000 | Figure S10B | GT |

**Table S2**: Decision cascades in the platelet: Efficient and robust but failing with aging

| |
|---|
| The platelet cascade is shown in Figure 2. Three features are combined here and support our considerations of decision processes as fast, robust heuristics versus exact mathematical decisions which may, however, sometimes take unforeseeably long to execute: |
| **I) Efficient platelet signaling cascade:** The platelet signaling cascade is fast and executes decisions very reliably according to all available evidence. Even multiple protein knockouts do not affect the central signaling process---it stays robust and fast. Concrete data that there is a central signaling backbone for efficient signaling transmission and that unless the central backbone is hit, otherwise processing is not harmed, both in mouse and man highly similar and well conserved signaling cascade and also for inhibitory signals (Balkenhol et al., 2020). In supplement, we show that TRPM7 KO affects signaling in model1 (without feedbacks and injury and healthy EC layer additions), but when feedbacks are involved, it does not change much because the signaling is more robust. |
| **II) Robust decision processes, decision funnels and activation control:** Platelet decision heuristics involve the SRC kinase and a bistability switch to ensure robust, fast, and direct decisions. This was confirmed by a combination of light-scattering-based thrombocyte aggregation data, western blot and calcium measurements (Mischnik et al., 2014). Evidence 2: Dynamic control centrality describes relaying functions as observed in signaling cascades of the SRC kinase (Karl and Dandekar, 2015b). The activating cascade is well conserved and robust, input from neighbor proteins modulates the signal, while the cytoskeleton executes the platelet activation as a subnetwork (Balkenhol *et al.*, 2020). |
| **III) However, shift in balance from fast, reliable execution by constant stress leads to thrombo-inflammation**<br>High stress, many activating inputs and inhibition signals at the same time lead to chronic inflammation, and too many cellular accidents due to aging result in chronic platelet pre-activation and thrombo-inflammation. This is almost a necessary consequence of the optimized cascade for blood-clotting in combination with infection protection, but with aging this leads to chronic inflammation: The rapid and robust decision due to positive feedbacks, parallel signaling pathways etc. works only well because it's controlled by self-brakes and EC control. When there is continuous activation from outside, although there is no injury, like in cancer and chronic inflammation (psoriasis, atherosclerosis, lupus etc.) and thrombo-inflammation (COVID-19, bacterial infections etc.), this leads to overactivation and trigger pathologies like thromboembolism etc. This effect is shown *in silico* in a simulation in the supplement. |

### Supplementary Figures

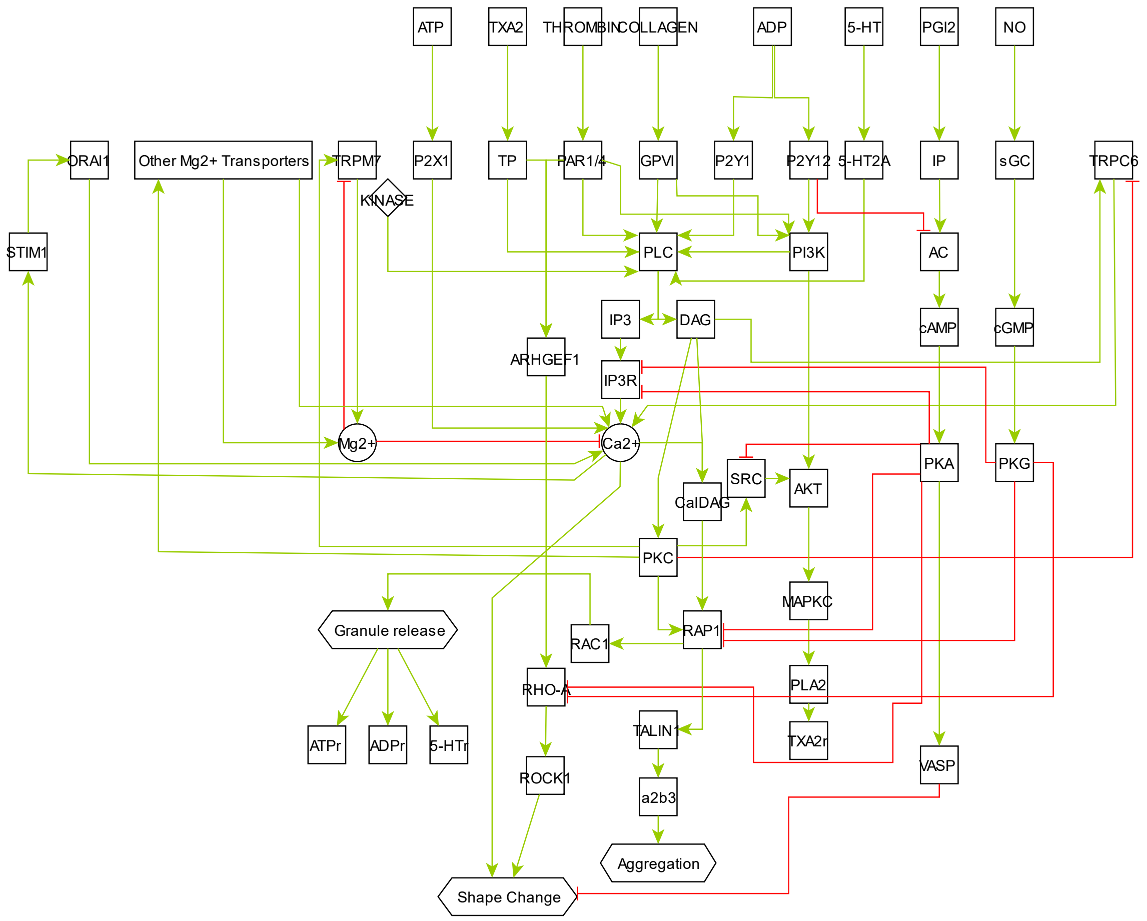


**Figure S1: Platelet signaling model without external control (representing *in vitro* conditions).** Topology of the underlying signaling network for simulations of in vitro experiments. The essential elements of the network are displayed in nodes, interconnected with either activating (green) or inhibiting (red) arrows. The difference from the model shown in the manuscript is the absence of amplification loops and internal and external inhibitory control.

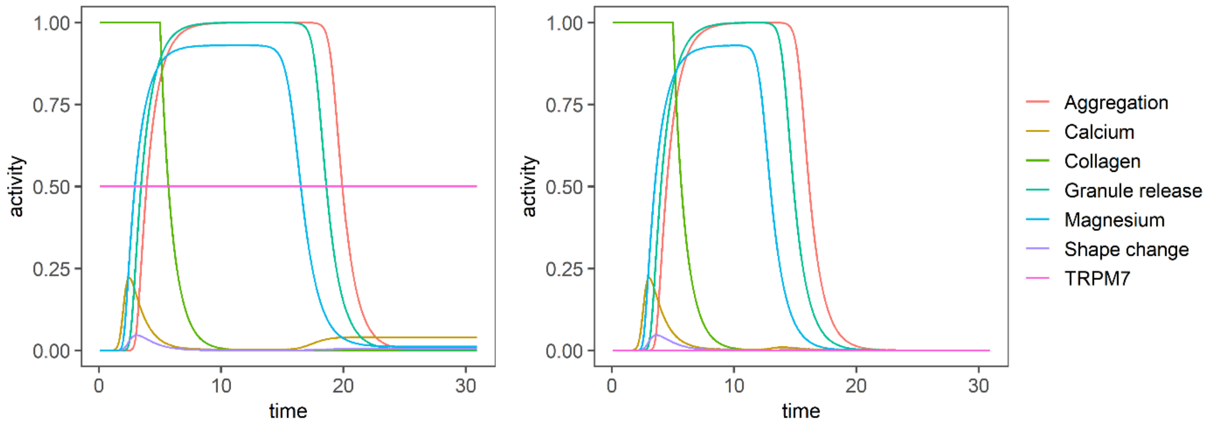


**Figure S2: Platelet signaling simulation without external control (representing in vitro conditions).** TRPM7 Kinase domain is set as constitutively active in wild-type (WT, left) and 0 in TRPM7 knockout (-/-, right). Activation of platelet cascade by collagen is simulated for two conditions of TRPM7: wild type (WT) in which TRPM7 is set to medium strength of 0.5 due to unknown kinetics of the kinase, and knockout (TRPM7$^{-/-}$) in which the kinase is set to 0. In both cases, the initial activation of PLC pathway leads to an increase in Ca2+ levels, and followingly, the ion channels for magnesium and calcium (e.g., TRPM7) are activated, balancing the Ca2+ levels with Mg2+, and the network then reaches the state of irreversible aggregation. When TRPM7 is inactivated (TRPM7$^{-/-}$), there is a slight delay in the activation of the cascade, and in the subsequent Ca2+ increase, the network stays active for a shorter duration.

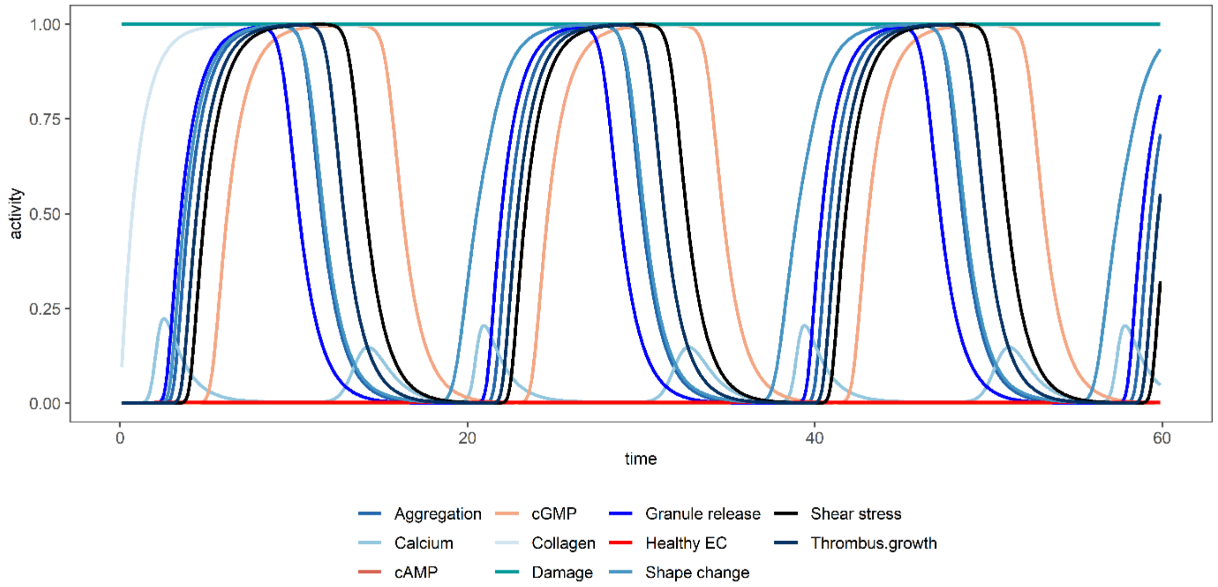


**Figure S3: Constant damage simulation.** Platelet activation persists, although some of the intrinsic self-inhibition mechanisms (e.g., RGS18-cGMP signaling) respond to platelet activation by trying to suppress it. However, continuous external input leads to repeated platelet activation (damage/injury: Value 1 from 0 to 1000).

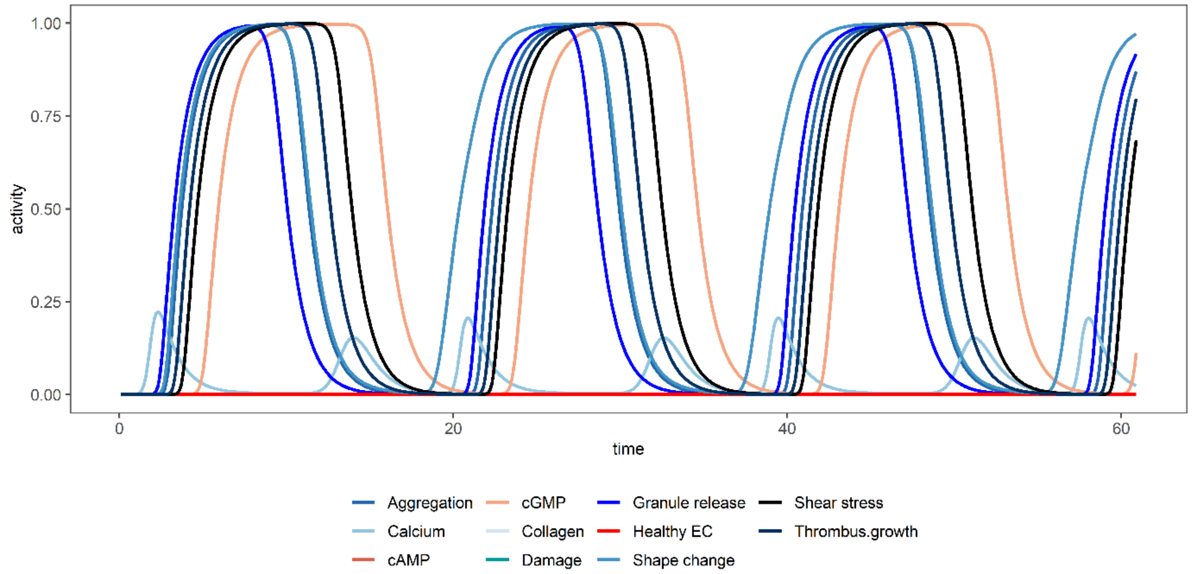


**Figure S4: EC absence simulation.** Platelet activation persists similarly to constant damage simulation, although some of the intrinsic self-inhibition mechanisms (e.g., RGS18-cGMP signaling) respond to platelet activation by trying to suppress it. However, lack of EC control leads to repeated platelet activation, even in the absence of damage (Healthy EC: Value 0 from 0 to 1000).

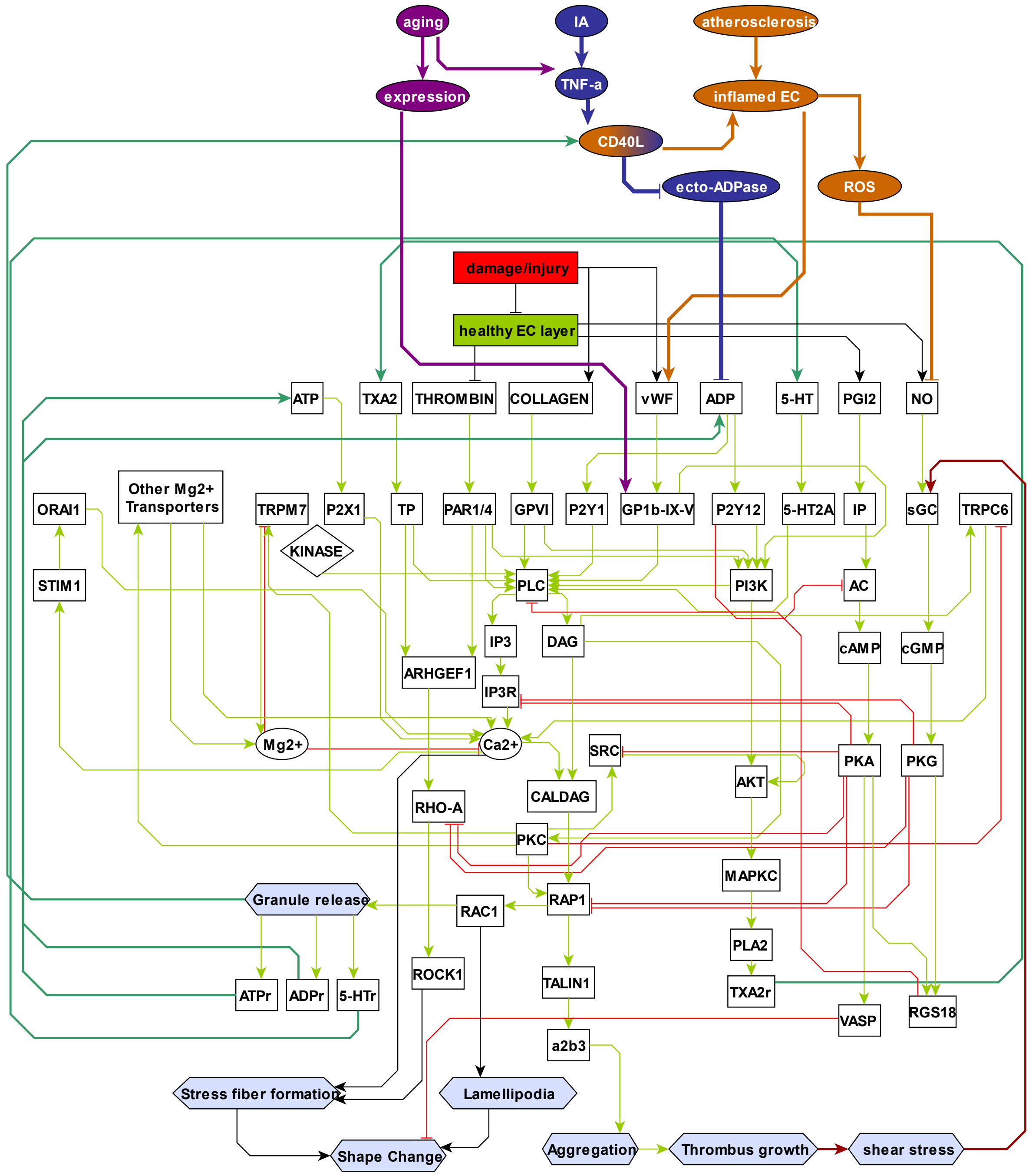


**Figure S5: Chronic inflammation diseases integrated in the model.** Effect of inflammation on platelet signaling is shown with a focus on atherosclerosis, inflammatory arthritis (IA), and aging. In atherosclerosis, endothelial cells (ECs) secrete vWF and promote platelet adhesion. In return, activated platelets release CD40L, which induces ECs to take on a more pro-inflammatory and pro-thrombotic profile, leading to more platelet activation, e.g., via inhibition of platelet-inhibitory cGMP signaling. In IA, ECs also show a pro-inflammatory and pro-thrombotic profile, and this increases platelet activation mainly via ADP signaling. Factors like CD40L inhibit the ecto-ADPase of ECs, promoting the ADP signaling even more. Lastly, in aging, although metabolic reprogramming is a main way of regulating platelets, we here only show the effects on signaling. This is exerted mainly via TNF-α secretion by monocytes and also via increased expression of platelet adhesion receptors, like GP1b-IX-V.

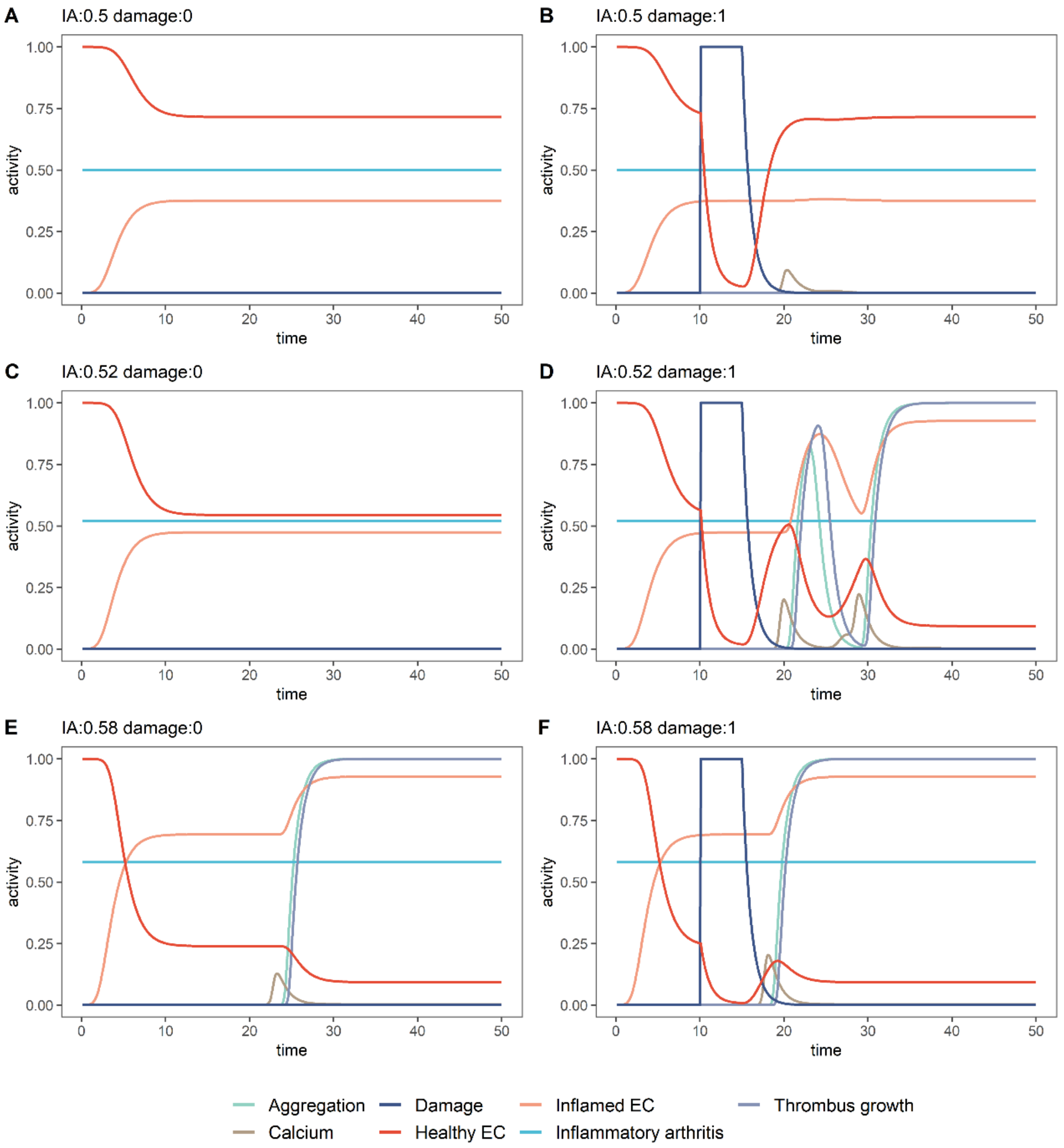


**Figure S6: Inflammatory arthritis (IA) simulation with damage input (right) and without damage (left).** Values below 0.52 for IA do not lead to any aberrations in platelet aggregation. Values between 0.52 and 0.58 only lead to aberrant aggregation when there is a damaged input. Values equal to or bigger than 0.58 lead to aberrant aggregation even though no damage is present.

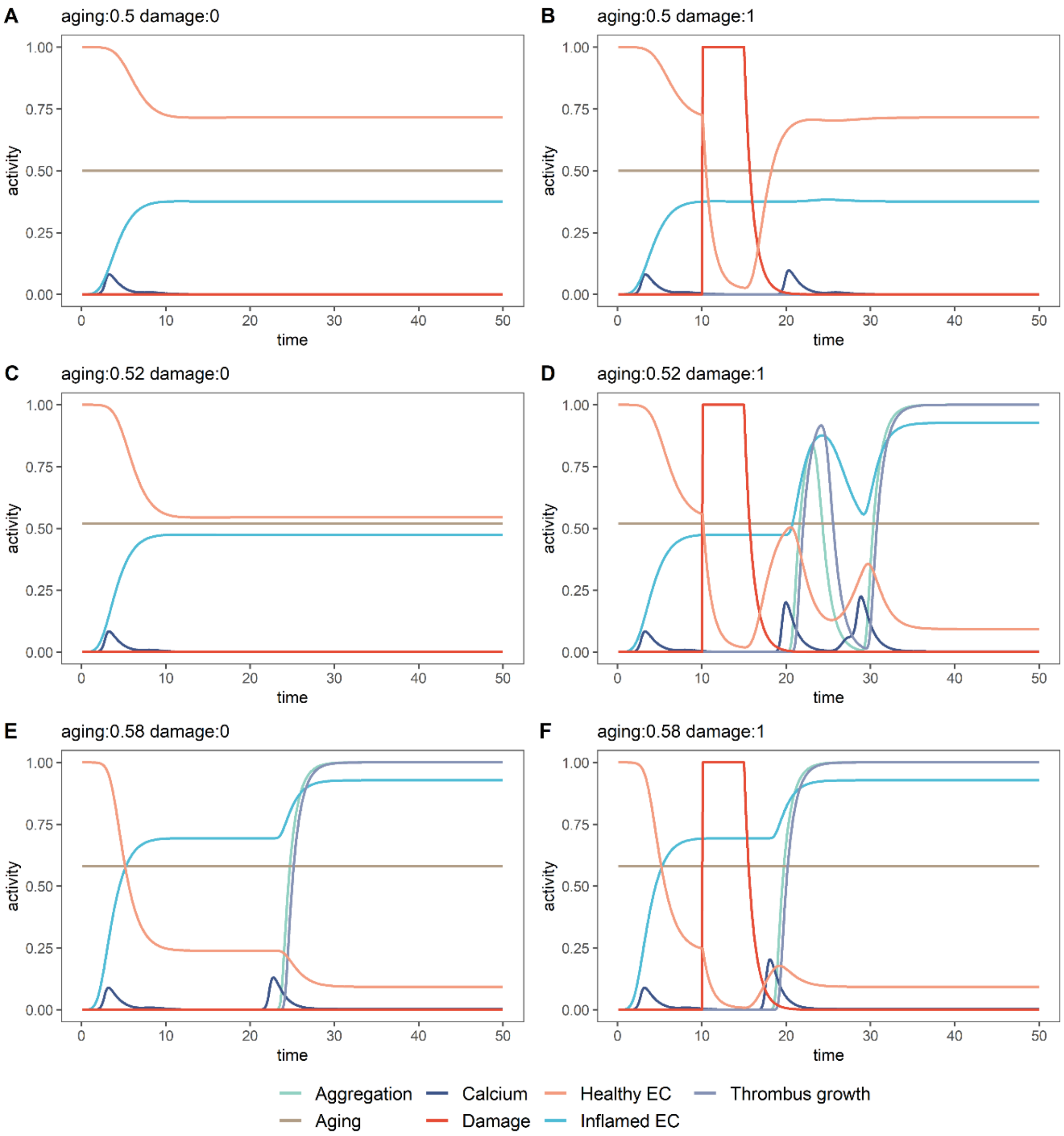


**Figure S7: Aging simulation with damage input (right) and without damage (left).** Values below 0.52 for aging do not lead to any aberrations in platelet aggregation. Values between 0.52 and 0.58 only lead to aberrant aggregation when there is a damaged input. Values equal to or bigger than 0.58 lead to aberrant aggregation even though no damage is present.

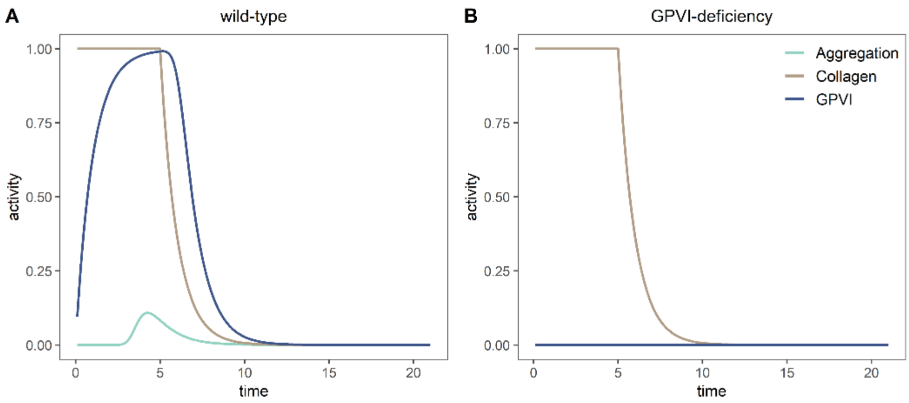


**Figure S8: Simulation of GPVI-deficiency (right) compared to wild-type simulations (left) in response to damage.** Platelet aggregation in response to collagen in wild-type platelets (A) is lost in GPVI deficiency (B).

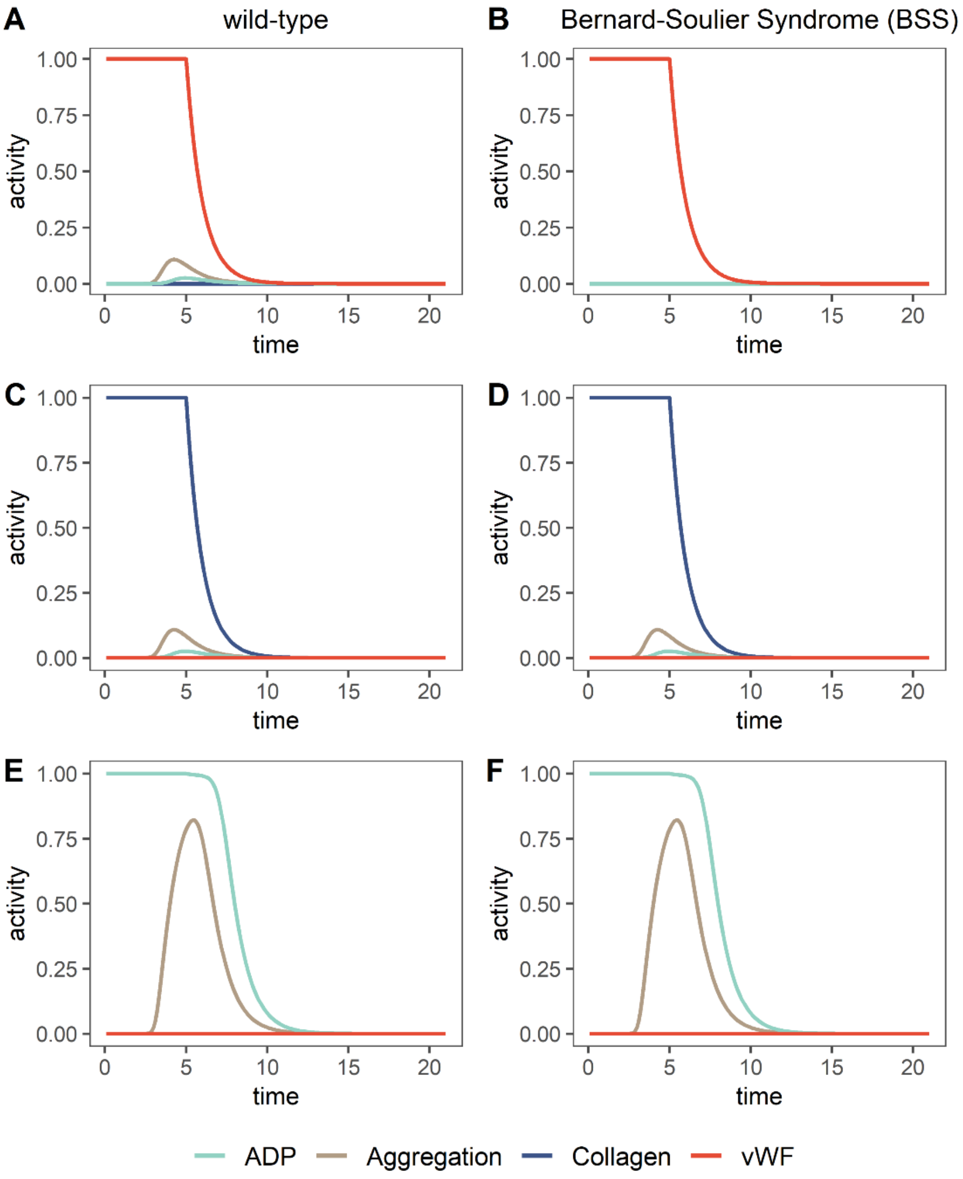


**Figure S9: Simulation of Bernard-Soulier Syndrome (BSS) (right) compared to wild-type simulations (left) in response to different agonists.** Platelet aggregation in response to vWF in wild-type platelets (A) is lost in BSS (B). However, other agonist-induced platelet aggregation responses are not affected (collagen (C-D), ADP (E-F)) in BSS.

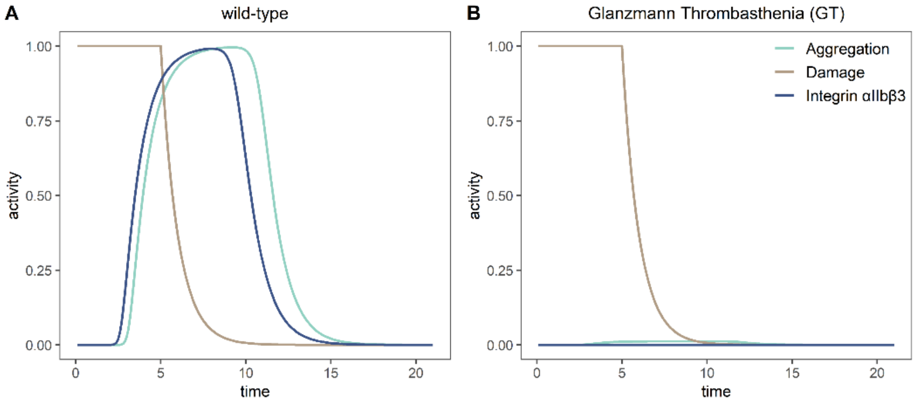


**Figure S10: Simulation of Glanzmann Thrombasthenia (GT) (right) compared to wild-type simulations (left) in response to damage.** Platelet aggregation in response to damage in wild-type platelets (A) is lost in GT (B).

### Supplementary Discussion

**Simulated network decision dynamics of platelet signalling agrees with physiology.**
When there is an injury, this state is rapidly transformed into platelet activation (**Fig. 4**). As a result of the injury, subendothelial collagen is exposed, and platelet inhibitory signaling from the healthy EC layer is interrupted until the damage is repaired. Here, initial activation arises from the binding of the collagen to the platelet receptors at the injury site (Stalker et al., 2012). This initial adhesion to collagen fibers is carried by the integrin α2β1 and the glycoprotein GPVI (ITAM signaling) (only GPVI is represented in **Fig. 4**) (Yeung et al., 2018). At high shear, binding of the GPIb-IX-V complex to vWF is also needed for stable adhesion (Estevez and Du, 2017). Adhesion initiates downstream signaling via PLCγ (denoted as PLC) and PI3Kβ (PI3K), leading to platelet activation (Muehlenkamp and Brausch, 2019; Yeung *et al.*, 2018). Platelet activation is followed by shape change, granule release (e.g., ADP from dense granules), and thromboxane2 (TXA2) generation via arachidonic acid (AA) cascade (Linkous and Yazlovitskaya, 2010). Furthermore, upon adhesion, thrombin is formed along the coagulation pathway via prothrombin (F2) on the surface of activated platelets (Yeung *et al.*, 2018).

Activation of PLC promotes an increase in the cytosolic calcium (Ca2+), activating Rap1B and RAC1, leading to cytoskeletal rearrangements involved in shape change. The platelets change from their natural discoid shape to a more extended shape via the formation of lamellipodia and filopodia. Similarly, Rac1 activation activates α-granule secretion (Golebiewska and Poole, 2013). Within α-granules, proplatelet agents like ADP, ATP, and serotonin (5-HT) are also released.

Granule release and TxA2 production leads to the amplification of the platelet activation signal by activating GPCRs (PAR1/4, P2Y1, P2Y12, P2X1, TP, 5-HT2A) and their downstream signaling through various G proteins. Thrombin receptor PAR1/4, ADP receptor P2Y1, TXA2 receptor TP, and serotonin receptor 5-HT2A act through $G\alpha_q$, which activates PLCβ. Gβγ is associated with receptors PAR1/4 and P2Y12 and activates PI3Kγ, while $G\alpha_{12/13}$ coupled to PAR1/4 and TP activates ARHGEF1 (followingly RHO-A). Moreover, $G\alpha_i$ coupled to P2Y12 inhibits AC and thus lifts platelet inhibition exerted by the cAMP signaling pathway (Li et al., 2010; Lopez-Vilchez et al., 2009; Nagy and Smolenski, 2018; Oury et al., 2015; Yeung *et al.*, 2018).

These signal transduction and amplification events during platelet activation then convert αIIbβ3 (a2b3 in the network) to its high-affinity active state for binding extracellular ligands such as fibrinogen or vWF. The αIIbβ3-fibrinogen or αIIbβ3-vWF complexes act as bridges between adjacent activated platelets and thus lead to platelet aggregation.

The positive feedback via granule release ensures a fast platelet aggregation that is needed in a time of injury. This robust mechanism (robustness against TRPM7 KO shown in **Fig. 5-6**) of rapidly recruiting more and more platelets ensures a fast response by amplifying the initial signal (Li *et al.*, 2010).

On the other hand, platelets do not aggregate without an end. The activation is terminated after the damage is repaired. This is achieved by various mechanisms, exemplified in our model by the negative feedback occurring in and outside the platelet. As the internal self-brake mechanism, we show RGS18, which is normally activated by platelet inhibitors (like cyclic nucleotides) and inhibits G protein signaling (Makhoul et al., 2018). In the network, we represent the internal mechanism by the inhibition that acts directly on PLC as a direct downstream of G protein signaling. The external self-brake also acts through cyclic nucleotides and functions through the mechanosensitive cGMP signaling. This mechanism includes the inhibition of thrombus growth by activation of guanylyl cyclase mechanically by the increasing shear stress in the bloodstream with the size of the clot formed (Feil et al., 2022).

**Decision funnels in platelets: effects of constant stimulation and chronic inflammation.**

The feedback loop via the external environment allows platelets to stabilize decision funnels or preferred decisions. The mechanisms for rapid decision indeed allow platelets to act rapidly, smoothly, and in a P-problem-like fashion: there is no stalling, and there is not much variation in decision time (in platelets measured by aggregometry). Hence, the platelets make a robust cellular decision. As a result, per time unit or biological system, there happens a surprisingly low amount of error. This also is expected to decrease the possible effect of a single mistake or mutation in a protein on the signaling outcome, which keeps hemostasis, an essential mechanism for survival, unaffected and functional. These single mistakes can have a more profound effect in the absence of signal amplification or the absence of the biological microenvironment (**Fig. 6 and S1-2**). On the other hand, under continuous stress, these robust and fast platelet decisions may lead to thrombo-inflammation because of constantly activating stimuli such as in diabetes, inflammation, infection, aging, or other conditions that increase oxidative stress and trigger various pathologies (**Fig. 7 and S5-7**, **Table S2**).

In thrombo-inflammation, the lack of EC control on thrombosis and inflammation leads to hyperactivation of platelets. As mentioned before, endothelial cells have an anti-thrombotic control over platelet function via various mechanisms. Likewise, ECs have an anti-inflammatory profile under healthy conditions. During thrombo-inflammation, inflammatory processes can shift the balance between pro- and anticoagulant properties of endothelium. This control can be lost, leading to continuous activation of platelets and recruitment of immune cells to the endothelium. Since platelets respond strongly to environmental stimuli, once the EC control is absent, they are hyperactive, and the environmental stimuli lead to their (self)-activating interaction with the endothelium and with various immune cells (Klavina et al., 2022; Sharma et al., 2022). Our simulations show recurring platelet activation in a loop in the continuous presence of procoagulant stimuli or the absence of EC control (**Fig. S3-4**).

- The role of platelets in chronic inflammation differs from the mechanisms in acute inflammation summarized above. In early phases of acute inflammation in response to bacterial or viral infection, platelet activation and coagulation help contain the pathogen dissemination at the site, partly by activating neutrophil NETosis (Stark and Massberg, 2021). In later stages, platelet activation inhibits neutrophils and suppresses TNF-α secretion from monocytes (Corken et al., 2014), leading to the resolution of the inflammation. The same receptor, GP1b, sustains both pro- and anti-inflammatory characteristics of platelets.

In chronic inflammation, the decision funnel switches in platelets from thrombosis and aggregation to favoring inflammation and thrombo-inflammation.

Hence, a change in platelets from pro-inflammatory to anti-inflammatory character is absent under these conditions (Corken *et al.*, 2014) as well as in ECs. The inflammation is associated with an increased risk of thrombosis (Stark and Massberg, 2021). More detailed literature comparison in the suppemental discussion.

**Shifting decision funnels and platelet system state in disease**

In atherosclerosis, which is a chronic inflammation disease, interaction of platelets with endothelial cells and immune cells promotes the atherosclerosis (Chen et al., 2020). Platelet adherence to the endothelium occurs through GP1bα and αIIbβ3 initiating the plaque formation (Stark and Massberg, 2021). Platelets further interact with the plaque via their GPVI receptors, earlier with laminin, and with collagen once the plaque raptures. GPVI signaling is essential in atherothrombosis and its inhibition was shown to inhibit thrombus formation (Jamasbi et al., 2015). Platelets do not contribute to the chronic inflammation in atherosclerosis only by their thrombotic function. They also recruit neutrophils and promote the adhesion of monocytes to the atherosclerotic endothelium and release cytokines and chemokines, increasing inflammation (such as CD154, IL1β, CCL5 (Klavina *et al.*, 2022), CXCL4, CXCL5, MIF (Stark and Massberg, 2021)). Platelets interaction with leukocytes and endothelial cells is mediated majorly by the platelet P-selectin (Klavina *et al.*, 2022) e.g., binding to PSGL1 on neutrophils, or interactions between platelet HMGB1 and neutrophil RAGE, platelet GP1bα and neutrophil integrin aMβ2 (MAC1) promoting neutrophil recruitment and NETosis. In return, neutrophils further activate platelets by the antimicrobial cathelicidin LL37/CRAMP they secrete via platelet GPVI signaling (Stark and Massberg, 2021).

What leads to artherothrombosis (plaque rupture) is continuous low-grade inflammation, during which ECs secrete vWF which promotes platelets' adhesion by their Gp1b receptor (GPIb-IV-X complex) activating platelet signaling. Platelet activation, in return, ends in release of factors like CD40 ligand, which induces ECs to produce more ROS, adhesion and inflammatory molecules such as IL1β, which induces ECs to express more adhesion proteins and promotes the adhesion of neutrophils and monocytes to the endothelium. At the same time, activated platelets also release PF-4 and more ROS. Production of ROS is observed to be driven via upregulating of NOX and COX-1, which then scavenges the NO in the environment, which normally is taken up by platelets and mediates inhibitory platelet signaling via cGMP. When the NO is scavenged by ROS, the absence of inhibitory signaling leads to more platelet activation and aggregation (Davi and Patrono, 2007). We modeled the effect of Ecs during the low-grade inflammation in atherosclerosis by adding the mechanisms summarized above (**Fig. 7**). We simulated different levels of atherosclerosis both in the absence of a damage/injury and when there is a damage stimulus. In all simulations, we started with a fully active "healthy EC layer" to simulate the changing balance between the healthy and inflamed EC layer. We introduced a damage stimulus at timepoint 10 to see the effects on a situation, where both inflammation and EC control is present at different levels. In the absence of damage, our simulations showed atherothrombosis only at high levels of inflammation caused by atherosclerosis (atherosclerosis $\geq$ 0.7). Medium levels of inflammation (0.52$\leq$atherosclerosis$<$0.7) led to aberrant platelet aggregation only in presence of a damage stimulus. In other words, in the medium range of atherosclerosis, a damage stimulus was enough to shift the balance from healthy EC to inflamed EC. However, low-grade inflammation (atherosclerosis $<$ 0.52) did not result in atherothrombosis even in presence of a damage stimulus, which failed to disrupt a healthy EC layer and its anti-thrombotic profile (**Fig. 7**).

Another chronic inflammatory disease, in which platelets show increased activity is inflammatory arthritis (IA). This activity is attributed to ADP signaling. ADP normally activates platelets by binding to P2Y12, which is a G-protein-coupled receptor. Downstream signaling leads to platelet activation and release of factors including CD40-ligand (CD40L), which is also induced by increased TNF-α levels in IA. CD40L, in return, induces Ecs to produce and secrete more pro-inflammatory factors and leads to the activation of more platelets by inhibiting the ecto-ADPase of endothelial cells resulting in higher amounts of available ADP in the bloodstream (Mac Mullan et al., 2010). We integrated the interplay between the IA-induced pro-thrombotic EC profile via ADP signaling and the platelet hyperactivation into our model (Fig. SÖ8) and simulated the effects of different levels of IA on platelet aggregation. We observed a similar trend as in atherosclerosis; however, IA exerted a stronger effect on platelet aggregation even at lower/medium levels (IA≥0.58) and ended up in full activation of platelet aggregation even in the absence of a damage input (**Fig. S6**).

High levels of TNF-α are also observed in the chronic inflammation during aging and is secreted by monocytes (Gu and Dayal, 2020). In return, activated platelets promote production of more cytokines by monocytes and also express surface proteins that increase the reactivity of platelets (Price et al., 2020). These platelets also showed higher surface expression of Gp1bα and GPIX compared to platelets from young mice (Gu and Dayal, 2020); they have in Fig. 6 a heatmap for scRNAseq data for platelet activation genes in old and young platelets). High levels of TNF-α asserts its effects by metabolic programming of the platelets, which leads to changes in pathways involved in mitochondrial processes. These effects of TNF-α were shown to be on thrombopoiesis and megakaryopoiesis rather than platelet signaling directly (Davizon-Castillo et al., 2019). We represented the effect of aging on expression of platelet receptors Gp1bα and GPIX and inflammation via TNF-α levels at the signaling level (metabolic reprogramming is not represented in our model) and our simulations showed almost identical results as platelet hyperactivation caused by IA-induced inflammation (**Fig. S7**).

***Decision funnel:*** *Platelet decisions happen in decision funnels, which helps model and understand platelet pathophysiology.* ***Hyperactivation***

Heuristic decision processes keep the platelets' response and cellular decision robust and fast by cascading activation of platelet signaling in the presence of external signals. Decision processes are not perfect, and decision funnels may shift under long-term stress, leading to disbalance in the platelet. Here, hyperfunctioning of platelets is more common than hypo-functioning, and particularly, the state of thrombo-inflammation is often reached.

Inherited platelet disorders easily trip the platelet signaling balance, majorly concerning the size of the platelets and their function. The first type of platelet disorder can lead to low counts of platelets (thrombocytopenia) or high counts of platelets (thrombocytosis) and is mostly caused by mutations in genes associated with megakaryopoiesis and thrombopoiesis. On the other hand, diseases concerning the platelet function are associated with hypofunctional platelets due to mutations in major platelet receptors like GPVI (**phenotype**: absent or severely reduced platelet aggregation in response to collagen and other GPVI agonists, confirmed/modeled by our simulations in **Fig. S8**), GP1b-GPIX-GPV (Bernard-Soulier Syndrome, **phenotype**: absence of ristocetin-induced platelet aggregation, reduced response to other agonists like collagen and ADP (Berndt and Andrews, 2011), reproduced by our simulations in **Fig. S9**), and αIIbβ3(Glanzmann Thrombasthenia (GT), phenotype: absent or severely reduced LTA with all agonists (ADP, TxA2, collagen, thrombin), confirmed by our simulations in **Fig. S10**) or in genes associated with platelet granules and signaling (Palma-Barqueros et al., 2021). However, inherited platelet disorders spanning 60 diseases are rare and vary from person to person in the clinical outcome (Bastida et al., 2019). A recent study showed a 3:$10^4$ prevalence for clinically relevant loss-of-function variants in platelet genes (Oved et al., 2021). On the other hand, thrombosis, caused by hyperfunctioning of the platelets, has been described as a major cause of death in the world, accounting for 25% of the deaths worldwide in 2010 (Wendelboe and Raskob, 2016). Thrombosis is the top cause of cardiovascular disorders like ischemic heart disease, stroke, and venous thromboembolism (VTE) (Raskob et al., 2014). Among the 42.0 million deaths from non-communicable diseases in 2019, ischemic heart disease caused 9.14 million and stroke caused 3.29 million deaths accounting for 29.5% of the deaths worldwide (Diseases and Injuries, 2020). Additionally, VTE was shown to have a yearly incidence rate of 1-2:1000 in Europe and the USA (Lutsey and Zakai, 2023). All combined account for a higher incidence rate than hypo-functioning platelet disorders, and there are far higher numbers of thrombotic events in addition to the deaths caused by the diseases listed, they are not sufficiently treated and not included in the incidence rates. We can confidently say that the hyperfunctioning of platelets is more common than hypo-functioning and this also is explained by how the platelet signaling is controlled to focusing on fast and cascading activation of platelet signaling in presence of external stimuli. This can also explain the weak effect of specific gene KOs on our model (except the receptors (**Fig. S8-10**)) in the presence of an external stimulus, e.g., damage to the endothelium. On the other hand, if we have a continuous stimulus or overactivation of a protein, the platelet activation status is irreversibly sustained at its maximum. This is what happens in pathologies involving hyperactivation of platelets, in which the sustained external stimulus or inhibited external EC control leads to aberrant activation of platelets. The second half of the cycle is closed by platelets activating the cause of their activation, be it inflammation or cancer.

**Table S3:** References for the interactions in the platelet model. To save space, the references are concisely given as Pubmed IDs (PMID).

| node1 | int | node2 | ref |
|---|---|---|---|
| **TRPM7** | + | Mg2+ | 29146750 |
| **TP** | + | ARHGEF1 | 14528298, 17298951 |
| **TP** | + | PLC | 29925522 |
| **PAR1/4** | + | PLC | 29925522 |
| **GPVI** | + | PLC | 29925522 |
| **P2Y1** | + | PLC | 29925522 |
| **PLC** | + | IP3 | 29925522, 29146750 |
| **PLC** | + | DAG | 29925522 |
| **PI3K** | + | AKT | 29925522, 30046761 |
| **SRC** | + | AKT | 23463387, 20586915 |
| **DAG** | + | CALDAG | 29925522 |

| | | | |
|---|---|---|---|
| **PKC** | + | SRC | 15314167 |
| **PKC** | + | RAP1 | 16979556 |
| **RAP1** | + | TALIN1 | 26929147, 35721481 |
| **RHO-A** | + | ROCK1 | 14660612, 23121917 |
| **P2Y12** | + | PI3K | 29925522 |
| **IP** | + | AC | 29925522, 30046761 |
| **AC** | + | cAMP | 29925522, 30046761 |
| **cAMP** | + | PKA | 28228483, 30046761 |
| **PKA** | + | VASP | 16197368 |
| **P2X1** | + | Ca2+ | 25709760 |
| **Mg2+** | - | Ca2+ | 29146750 |
| **PKA** | - | SRC | 23463387, 12606547 |
| **PKA** | - | RAP1 | 26929147 |
| **P2Y12** | - | AC | 29925522 |
| **PAR1/4** | + | ARHGEF1 | 29925522, 17298951 |
| **PKA** | - | IP3R | 26929147, 30046761 |
| **PKA** | - | RHO-A | 24100445, 30046761 |
| **ATP** | + | P2X1 | 25709760 |
| **TXA2** | + | TP | 29925522 |
| **THROMBIN** | + | PAR1/4 | 29925522 |
| **COLLAGEN** | + | GPVI | 29925522 |
| **ADP** | + | P2Y1 | 29925522 |
| **ADP** | + | P2Y12 | 29925522 |
| **PGI2** | + | IP | 29925522 |
| **Ca2+** | + | CALDAG | 29925522 |
| **PAR1/4** | + | PI3K | 29925522 |
| **5-HT** | + | 5-HT2A | 19541671, 21071698 |
| **5-HT2A** | + | PLC | 21071698 |
| **KINASE** | + | PLC | 29146750, 22759789 |
| **Ca2+** | + | STIM1 | 29146750, 16582901, 18559454 |
| **DAG** | + | TRPC6 | 9930701 |
| **TRPC6** | + | Ca2+ | 9930701 |
| **ORAI1** | + | Ca2+ | 29146750 |
| **PKC** | + | Other Mg2+ Transporters | 23727583, 15804357 |
| **PKC** | + | TRPM7 | there is evidence for PKA --> TRPM7 in 15069188 |
| **PKC** | - | TRPC6 | 20961851, 16772730, 11278449, 12721302 |
| **a2b3** | + | Aggregation | 29550521, 15696195 |
| **RAP1** | + | RAC1 | 24588354 |
| **Other Mg2+ Transporters** | + | Mg2+ | 15804357 |
| **Granule release** | + | ADPr | 29925522, 24588354 |
| **Granule release** | + | ATPr | 29925522, 24588354 |
| **Granule release** | + | 5-HTr | 29925522, 24588354 |
| **VASP** | - | Shape Change | 16197368, 26929147 |
| **MAPKC** | + | PLA2 | 20642808, 23620575 |
| **PLA2** | + | TXA2r | 20642808 |
| **Mg2+** | - | TRPM7 | 29500477, 17217066, 11385574, 22759789 |
| **Other Mg2+ Transporters** | + | Ca2+ | |
| **IP3** | + | IP3R | 29925522, 28228483, 29146750 |
| **IP3R** | + | Ca2+ | 29925522, 28228483, 29146750 |
| **sGC** | + | cGMP | 20929147, 30046761 |
| **cGMP** | + | PKG | 20929147 |
| **PKG** | - | RAP1 | 20929147 |
| **PKG** | - | IP3R | 20929147 |
| **PI3K** | + | PLC | hsa04611 |
| **GPVI** | + | PI3K | 30429532 |
| **AKT** | + | MAPKC | 29925522, 28228483 |
| **DAG** | + | PKC | 29925522 |

| | | | |
|---|---|---|---|
| **CALDAG** | + | RAP1 | 29925522, 26929147 |
| **NO** | + | sGC | 26929147, 30046761 |
| **STIM1** | + | ORAI1 | 29146750, 16582901, 18559454 |
| **ARHGEF1** | + | RHO-A | 14528298, 17298951 |
| **TALIN1** | + | a2b3 | 26929147, 35721481 |
| **RAC1** | + | Granule release | 24588354 |
| **ADPr** | + | ADP | released |
| **ATPr** | + | ATP | released |
| **TXA2r** | + | TXA2 | released |
| **5-HTr** | | 5-HT | released |
| **PKG** | - | RHO-A | 29550521 |
| **Aggregation** | + | Thrombus growth | 30327468 |
| **Thrombus growth** | + | shear stress | 30327468 |
| **shear stress** | + | sGC | 30327468 |
| **PKA** | | RGS18 | 30046761 |
| **PKG** | | RGS18 | 30046761 |
| **RGS18** | - | PLC | 30046761 |
| **healthy EC layer** | - | THROMBIN | 26481314, 30429532 |
| **healthy EC layer** | + | PGI2 | 26481314 |
| **healthy EC layer** | + | NO | 26481314 |
| **damage/injury** | - | healthy EC layer | logic |
| **damage/injury** | + | COLLAGEN | 29925522, 29340130 |
| **RAC1** | + | Lamellipodia | 23121917, 16195235, 12453877 |
| **ROCK1** | + | Stress fiber formation | 23121917 |
| **Ca2+** | + | Stress fiber formation | 23121917 |
| **Stress fiber formation** | + | Shape Change | 23121917 |
| **Lamellipodia** | + | Shape Change | 23121917 |
| **GP1b-IX-V** | + | PI3K | 35615944 |
| **GP1b-IX-V** | + | PLC | 35615944 |
| **vWF** | + | GP1b-IX-V | 28228483, 35615944 |
| **damage/injury** | + | vWF | 28228483, 35615944 |
